# Circadian control of intrinsic heart rate via a sinus node clock and the pacemaker channel

**DOI:** 10.1101/684209

**Authors:** Yanwen Wang, Servé Olieslagers, Anne Berit Johnsen, Svetlana Mastitskaya, Haibo Ni, Yu Zhang, Nicholas Black, Cali Anderson, Charlotte Cox, Annalisa Bucchi, Sven Wegner, Beatriz Bano-Otalora, Cheryl Petit, Eleanor Gill, Sunil Jit Logantha, Nick Ashton, George Hart, Henggui Zhang, Elizabeth Cartwright, Ulrik Wisloff, Paula Da Costa Martins, Dario DiFrancesco, Halina Dobrzynski, Hugh D. Piggins, Mark R. Boyett, Alicia D’Souza

## Abstract

In the human, there is a circadian rhythm in the resting heart rate and it is higher during the day in preparation for physical activity. Conversely, slow heart rhythms (bradyarrhythmias) occur primarily at night. Although the lower heart rate at night is widely assumed to be neural in origin (the result of high vagal tone), the objective of the study was to test whether there is an *intrinsic* change in heart rate driven by a local circadian clock. In the mouse, there was a circadian rhythm in the heart rate *in vivo* in the conscious telemetrized animal, but there was also a circadian rhythm in the intrinsic heart rate in denervated preparations: the Langendorff-perfused heart and isolated sinus node. In the sinus node, experiments (qPCR and bioluminescence recordings in mice with a *Per1* luciferase reporter) revealed functioning canonical clock genes, e.g. *Bmal1* and *Per1*. We identified a circadian rhythm in the expression of key ion channels, notably the pacemaker channel *Hcn4* (mRNA and protein) and the corresponding ionic current (funny current, measured by whole cell patch clamp in isolated sinus node cells). Block of funny current in the isolated sinus node abolished the circadian rhythm in the intrinsic heart rate. Incapacitating the local clock (by cardiac-specific knockout of *Bmal1*) abolished the normal circadian rhythm of *Hcn4*, funny current and the intrinsic heart rate. Chromatin immunoprecipitation demonstrated that *Hcn4* is a transcriptional target of BMAL1 establishing a pathway by which the local clock can regulate heart rate. In conclusion, there is a circadian rhythm in the intrinsic heart rate as a result of a local circadian clock in the sinus node that drives rhythmic expression of *Hcn4*. The data reveal a novel regulator of heart rate and mechanistic insight into the occurrence of bradyarrhythmias at night.

---

Living things including humans are attuned to the 24 h day-night cycle and many biological processes exhibit an intrinsic ~24 h (i.e. circadian) rhythm. In the human, the resting heart rate (in the absence of physical activity) shows a circadian rhythm and is higher during the day when we are awake^1,2^. The heart is therefore primed, anticipating the increase in demand during the awake period. Conversely, bradyarrhythmias primarily occur at night^1,3^. Previously, the circadian rhythm in heart rate *in vivo* has been attributed to the autonomic nervous system: to high sympathetic nerve activity accompanying physical activity during the awake period and high vagal tone during the sleep period^4^. This is partly based on heart rate variability as a surrogate measure of autonomic tone^5–7^; however, we have shown that heart rate variability cannot be used in any simple way as a measure of autonomic tone^8^. Therefore, the mechanism underlying the circadian rhythm in heart rate is still unknown. We asked the question whether there is a circadian rhythm in the *intrinsic* heart rate set by the pacemaker of the heart, the sinus node.

In a cohort of nine mice, ECG telemetry *in vivo* showed a circadian rhythm in mean heart rate and other electrophysiological parameters (PR interval, QRS duration and QT interval) during a 12 h light:12 h dark lighting regime, and circadian rhythms in these parameters were sustained when the mice were placed in constant dark conditions (Figure 1). A circadian rhythm in heart rate, PR interval, QRS duration and QT interval are also observed in the human^7,9–11^. Mice are nocturnal and as expected they rested more and were less active from Zeitgeber time (ZT) 0 to ZT 12 (lights-on) and were more active from ZT 12 to ZT 0 (lights-off); this activity pattern continued in constant darkness (Figure 1)^12,13^. The heart rate was highest at ~ZT 13 and it varied by 76±4 beats/min (as determined by the least-squares best fit sine wave) over the course of 24 h (Figure 1; Table S1 in the Supplementary Information).

**Figure 1.**
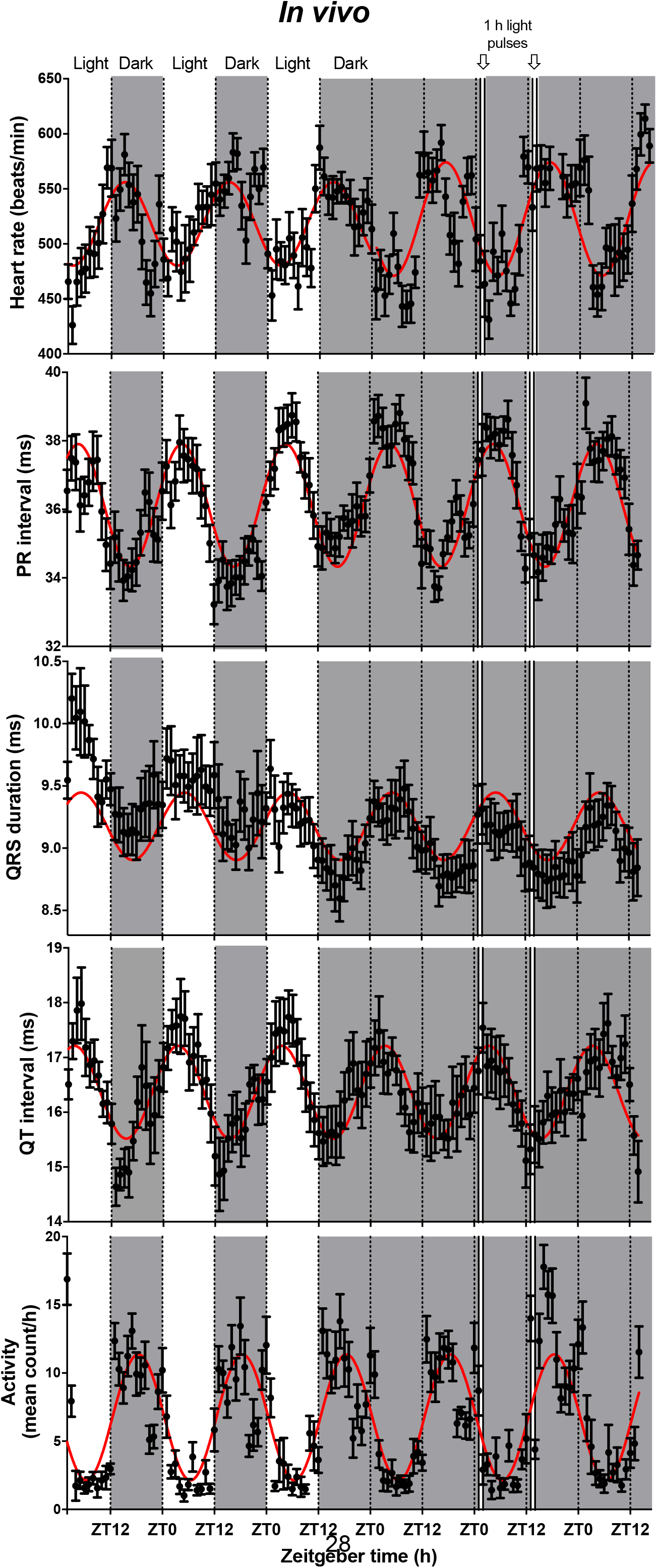
Circadian rhythm in heart rate and other ECG parameters. Circadian rhythm of heart rate, PR interval, QRS duration, QT interval and physical activity (measured using telemetry) in conscious mice (n=9) over ~6 days. Light and shaded regions represent light and dark phases in this and all similar figures – there was a 12 h light/12 h dark cycle for the first three days and constant darkness subsequently. Timing of 1 h light pulses shown. In this and all similar figures, means±SEM are shown and data are fit with a standard sine wave shown in red.

## Circadian rhythm in intrinsic sinus node pacemaking and ion channel expression

Experiments on the isolated, denervated, Langendorff-perfused heart (Figure 2A) dissected at projected ZT 0 or ZT 12 demonstrated that there was a circadian rhythm in *intrinsic* sinus node pacemaking: the intrinsic heart rate was still higher at ZT 12 than ZT 0 – by 107 beats/min (Table S1). In the Langendorff-perfused heart, a suitable pacing protocol (Figure S1 in the Supplementary Information) showed the corrected sinus node recovery time (cSNRT), a commonly used indicator of sinus node function, to be significantly different at ZT 0 and ZT 12 (Figure 2B). It is concluded that there is a circadian rhythm in the intrinsic heart rate. What is responsible for the intrinsic circadian rhythm? Molecular circadian clock machinery is ultimately responsible for all circadian rhythms. The master clock is in the suprachiasmatic nucleus of the brain, but there are peripheral clocks in other organs^14^. To determine if there is circadian clock machinery in the sinus node, qPCR was used – qPCR showed that mRNA for a wide range of molecules known to be involved in the circadian clock is present in the sinus node and the expression varied in the expected manner from ZT 0 to ZT 12 (Figure 2C). Two key transcripts, *Bmal1* and *Clock*, were measured in the sinus node at four time points (BMAL1 and CLOCK form a heterodimer); Figure 2D,E shows that they varied in a circadian manner and were at a maximum at ~ZT 0 (Table S1). The presence of an intrinsic circadian clock in the sinus node was confirmed by measuring the bioluminescence in the isolated sinus node from a transgenic mouse (*Per1*∷LUC) carrying a luciferase reporter gene reporting the activity of *Per1*, a key circadian clock component (Figure 2F). *Per1*-driven bioluminescence fluctuated in the expected circadian manner and this periodicity was lost in *Cry1*^−/−^ *Cry2*^−/−^ mice lacking the *Cryptochrome* genes and consequently lacking a functional clock^15^ (Figure 2F). Pacemaking is the result of the concerted action of ion channels and Ca^2+^-handling proteins and it is likely that the local circadian clock in the sinus node controls the heart rate by controlling their expression. Expression of mRNA for many of these molecules (and also some key regulatory transcription factors) was measured by qPCR and some transcripts, for example the pacemaking ion channel *Hcn4*, demonstrated significant daily rhythms (Figure 2G; Table S2).

**Figure 2.**
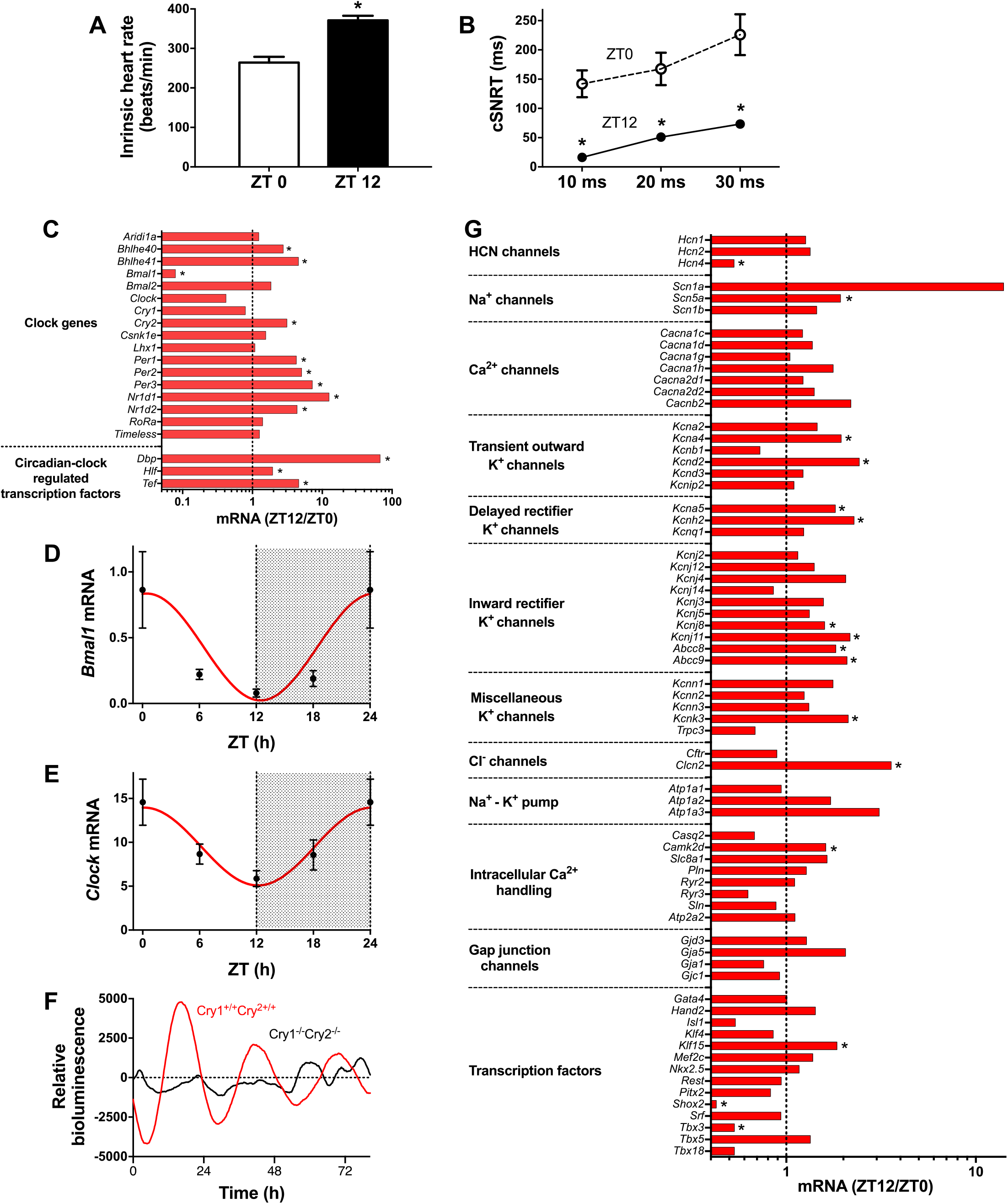
Circadian rhythm in intrinsic heart rate, local circadian clock in sinus node, and circadian rhythm in expression of ion channels and other molecules underlying pacemaker activity of sinus node. **A**, Intrinsic heart rate measured from the Langendorff-perfused heart isolated at ZT 0 and ZT 12 (n=9 and 8 mice). *P<0.05; two-tailed unpaired t test with Welch’s correction. **B**, Corrected sinus node recovery time (cSNRT) at three pacing cycle lengths measured from the Langendorff-perfused heart isolated at ZT 0 and ZT 12 (n=9 and 8 mice). *ZT 12 versus ZT 0, P<0.05; two-way ANOVA with Holm-Šídák post-hoc test. **C**, Relative expression of transcripts encoding key clock components in the sinus node at ZT 12 (as compared to ZT 0; n=7 and 9 mice). The vertical line corresponds to 1, i.e. no change. Values <1 correspond to a decrease at ZT 12 and >1 an increase. *ZT 12 versus ZT 0, P<0.05; limma test followed by Benjamini-Hochberg False Discovery Rate correction (set at 5%). **D** and **E**, Expression of *Bmal1* and *Clock* mRNA in the sinus node at four time points over 24 h (n=6 mice for ZT 0; n=6 for ZT 6; n=7 for ZT 12; n=8 for ZT 18). **F**, *Per1* activity (reported by luciferase bioluminescence) in the isolated sinus node from *Per1*∷LUC mice on a *Cry1*^+/+^*Cry2*^+/+^ or *Cry1*^−/−^*Cry2*^−/−^ background. Similar data obtained from two other *Cry1*^+/+^*Cry2*^+/+^ mice and two other *Cry1*^−/−^*Cry2*^−/−^ mice. **G**, Relative expression of transcripts encoding ion channels, Na^+^-K^+^ pump subunits, intracellular Ca^2+^-handling molecules, gap junction channels and key transcription factors in the sinus node at ZT 12 (as compared to ZT 0; n=7 and 9 mice). *ZT 12 versus ZT 0, P<0.05; Limma test followed by Benjamini-Hochberg False Discovery Rate correction (set at 5%).

## Circadian rhythm in HCN4 and *I*_f_

Of the potential mechanisms controlling the intrinsic heart rate, we principally focused on *Hcn4*, because this is known to play a central role in pacemaking in the sinus node^16^. *Hcn4* mRNA was measured at six time points and was at a maximum at ~ZT 0 (Figure 3A; Table S1). Expression of HCN4 protein in the sinus node was measured using western blot at ZT 0 and ZT 12 – the western blot is shown in Figure 3B and the mean HCN4 band intensities are shown in Figure 3C. HCN4 protein expression was higher at ZT 12 than ZT 0 (Figure 3C). Expression of HCN4 protein in the sinus node was also measured using immunohistochemistry (Figure 3D,E). Figure 3D shows examples of immunolabelling of HCN4 in tissue sections through the sinus node from mice culled at ZT 0 and ZT 12); consistent with the western blot, the labelling was brighter, indicating higher expression, at ZT 12. This is confirmed by the mean data in Figure 3E. In summary, the data show a daily rhythm in HCN4 protein, but it is out of phase with the circadian rhythm in mRNA. Western blot and immunohistochemistry were used to measure HCN4 at four time points and this showed that HCN4 protein peaked at between ZT 6 and 12 (Figure S2; Table S1) – HCN4 protein, therefore, lags behind mRNA - this is expected. If there are changes in HCN4 protein, there should be changes in the corresponding ionic current, *I*_f_. Figure 4A shows patch clamp recordings of *I*_f_ from isolated sinus node cells – the cells were isolated at ZT 0 and ZT 10 and the recordings were made at ZT 2 and ZT 12 (the earliest time points at which recordings could be made). The current density was approximately double at ZT12 than at ZT2 and this is confirmed by the mean current-voltage relationships for *I*_f_ at the two time points in Figure 4B. Once cells were isolated at ZT 0 and ZT 10, recordings from different cells could continue to be made for ~3 h. In the case of cells isolated at ZT 0 we observed the current density to increase the later the recording, but in the case of cells isolated at ZT 10 we observed the current density to decrease the later the recording. In Figure 4C we have plotted the density of *I*_f_ in a total of 367 sinus node cells against the time of the recording (this includes cells which were isolated at ZT 3, 6, 9, 12 and 15). Figure 4C shows a projected daily rhythm in the density of *I*_f_ – clearly the circadian clock does not stop ticking on isolation of single sinus node cells. *I*_f_ density reached a maximum at ~ZT 10, approximately at the same time as the peak in HCN4 protein as assessed by western blot and immunohistochemistry (Figure S2; Table S1). Current density determines the rate of change of membrane potential and is, therefore, the physiologically important variable. It is calculated by dividing the current amplitude by the cell capacitance, C_m_, and these variables are shown in Figure 4D,E. As anticipated there is a projected daily rhythm in *I*_f_ amplitude, which peaks at the same time as *I*_f_ density, but the fractional change in amplitude is less than the fractional change of density (Figure 4D,E). This is because there is a circadian rhythm in C_m_ (Figure 4E); in other words, a circadian rhythm in C_m_ contributes to the circadian rhythm in *I*_f_ density. C_m_ is determined by cell size and other experiments confirmed that there is a circadian rhythm in cell size (Figure S3). A circadian rhythm in cell size may be surprising. However, a circadian rhythm in cell size has been reported in other tissues: in mouse liver, cell size at ZT 12 is less than at ZT 2^17^; axonal volume shows a circadian rhythm in s-LNv clock neurons in Drosophila^18^; glial cells and neurons in the housefly’s visual system show rhythmic size changes^19^.

**Figure 3.**
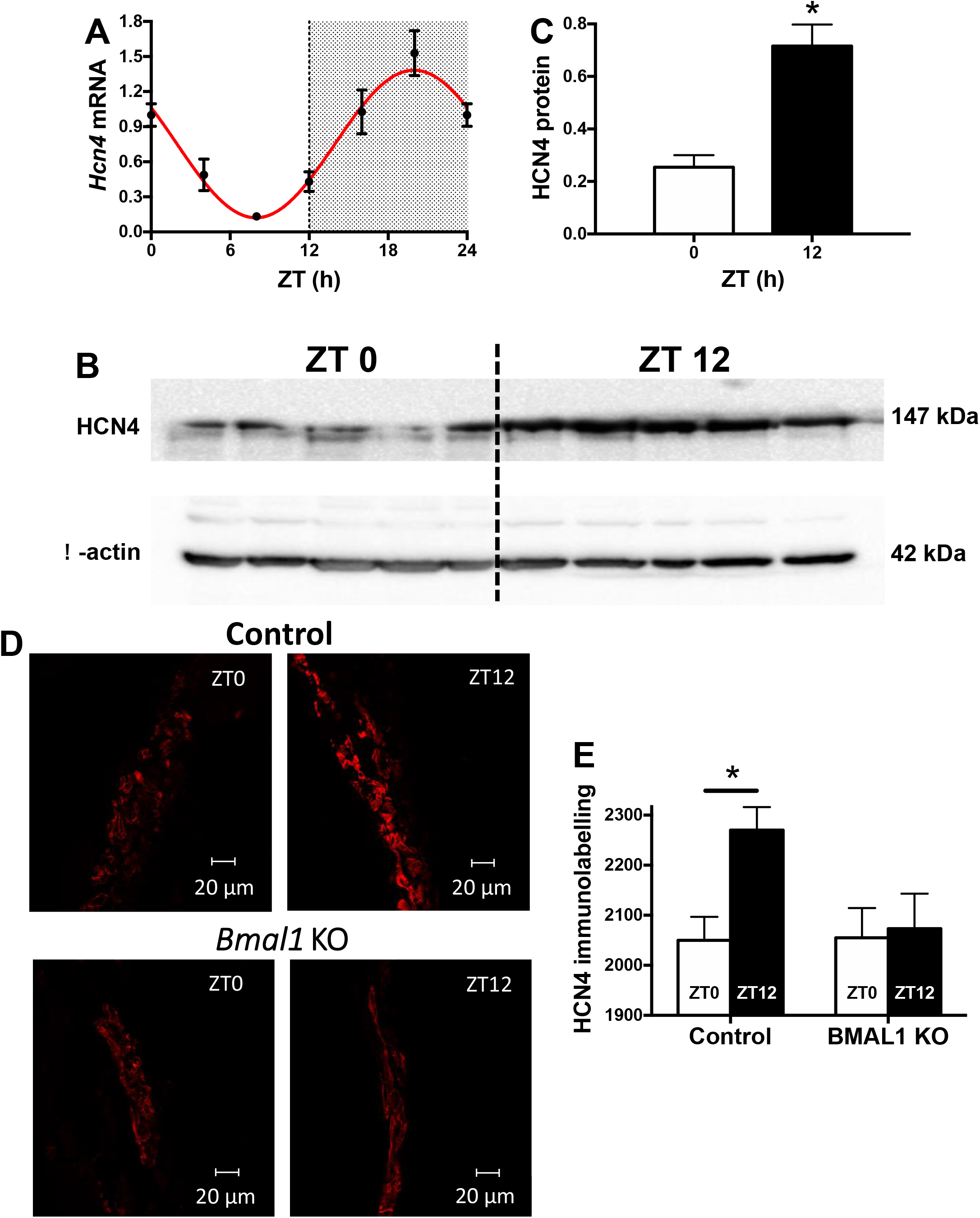
Diurnal rhythm in *Hcn4* mRNA and protein in sinus node. **A**, Expression of *Hcn4* mRNA in the sinus node at six time points over 24 h (n=6 mice for ZT 0; n=6 for ZT 4; n=7 for ZT 8; n=7 for ZT 12; n=7 for ZT 16; n=5 for ZT 20). **B**, Western blot for HCN4 and the housekeeper, β-actin, in the sinus node of 5 mice culled at ZT 0 and 5 mice culled at ZT 12. **C**, Mean HCN4 protein expression (relative to the expression of the housekeeper) in the sinus node determined by western blot at ZT 0 and ZT 12. *P<0.05; two-tailed unpaired t test with Welch’s correction. **D**, Immunolabelling of HCN4 protein (red signal) in sections through the sinus node dissected from control wild-type (top) and cardiac-specific *Bmal1* knockout mice culled at ZT 0 and ZT 12. **E**, Mean HCN4 protein expression determined by immunohistochemistry in the sinus node of control wild-type and cardiac-specific *Bmal1* knockout mice at ZT 0 and ZT 12 (wild-type mice, n=56/42 sections from 3/3 mice; cardiac-specific *Bmal1* knockout mice, n=38/46 sections from 3/3 mice). *P<0.05; two-way ANOVA with Holm-Šídák post-hoc test.

**Figure 4.**
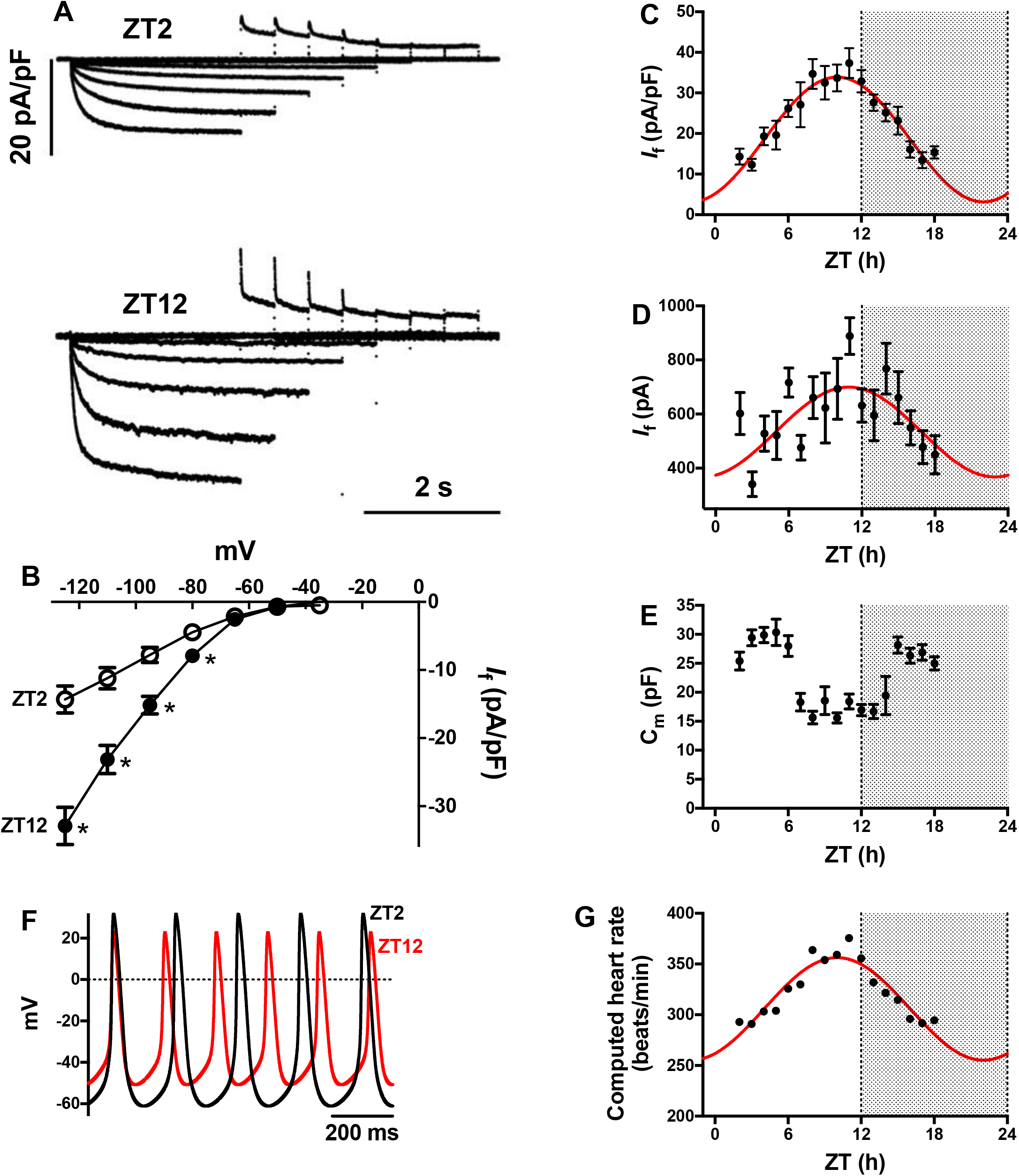
Circadian rhythm in *I*_f_ in sinus node. **A**, Families of recordings of *I*_f_ from sinus node cells made at ZT 2 and ZT 12. **B**, Current-voltage relationships for *I*_f_ recorded at ZT 2 (n=10 cells/3 mice) and ZT 12 (n=16 cells/3 mice). *ZT 12 versus ZT 0, P<0.05; two-way ANOVA with Holm-Šídák post-hoc test. **C**, Density of *I*_f_ at −125 mV over 24 h (n=10 cells for ZT 2; n=11 for ZT 3; n=10 for ZT 4; n=12 for ZT 5; n=8 for ZT 6; n=16 for ZT 7; n=14 for ZT 8; n=14 for ZT 9; n=27 for ZT 10; n=16 for ZT 11; n=16 for ZT 12; n=18 for ZT 13; n=13 for ZT 14; n=15 for ZT 15; n=17 for ZT 16; n=16 for ZT 17; n=11 for ZT 18; from 24 mice); *I*_f_ density is plotted against the time of recording. **D**, Amplitude of *I*_f_ at −125 mV over 24 h (from same experiments as for C). **E**, C_m_ over 24 h (from same experiments as for C). **F**, Computed mouse sinus node action potentials at ZT 2 and ZT 12 based on a model incorporating the measured differences in the density of *I*_f_. **G**, Computed ‘heart rate’ of a mouse sinus node cell over 24 h based on a model incorporating the measured differences in density of *I*_f_.

## Role of *I*_f_ and other ionic mechanisms in circadian rhythm in intrinsic heart rate

To test whether the changes in *I*_f_ density could account for the daily rhythm in the intrinsic heart rate, the observed changes in *I*_f_ density (Figure 4C) were incorporated into a biophysical model of the spontaneous action potential of the mouse sinus node (Figure 4F). The model predicted a circadian rhythm in heart rate of 101±11 beats/min with a maximum heart rate at ~ZT 10 (Figure 4G; Table S1), roughly consistent with experimental data for the intrinsic heart rate (Figure 2A). This suggests that HCN4 and *I*_f_ participate in the circadian rhythm in the intrinsic heart rate. This was confirmed in the isolated, denervated sinus node dissected at four different time points. Block of *I*_f_ by 2 mM Cs^+^ (selective blocker of *I*_f_^20^) decreased the heart rate as expected (Fig. 5A, top), but the effect of Cs^+^ on heart rate was greater at ZT 12 than ZT 0 (Figure 5A, bottom). Furthermore, in the presence of Cs^+^, the circadian rhythm in the intrinsic heart rate was abolished (Figure 5A, top). Analogous results were obtained *in vivo*. In the conscious mouse, the heart rate at ZT 0 and ZT 12 was measured using an ECGenie (a platform with embedded ECG electrodes). An intraperitoneal injection of 6 mg/kg ivabradine (selective blocker of *I*_f_^21^) was given to block *I*_f_. Once again, block of *I*_f_ decreased the heart rate as expected (Figure 5C) and the effect of ivabradine on heart rate was greater at ZT 12 than ZT 0 (Figure 5D). Furthermore, in the presence of ivabradine, the circadian rhythm in the heart rate *in vivo* was abolished (Figure 5C). It is concluded that HCN4 and *I*_f_ participate in the circadian rhythm in the intrinsic heart rate as well as the heart rate *in vivo*.

**Figure 5.**
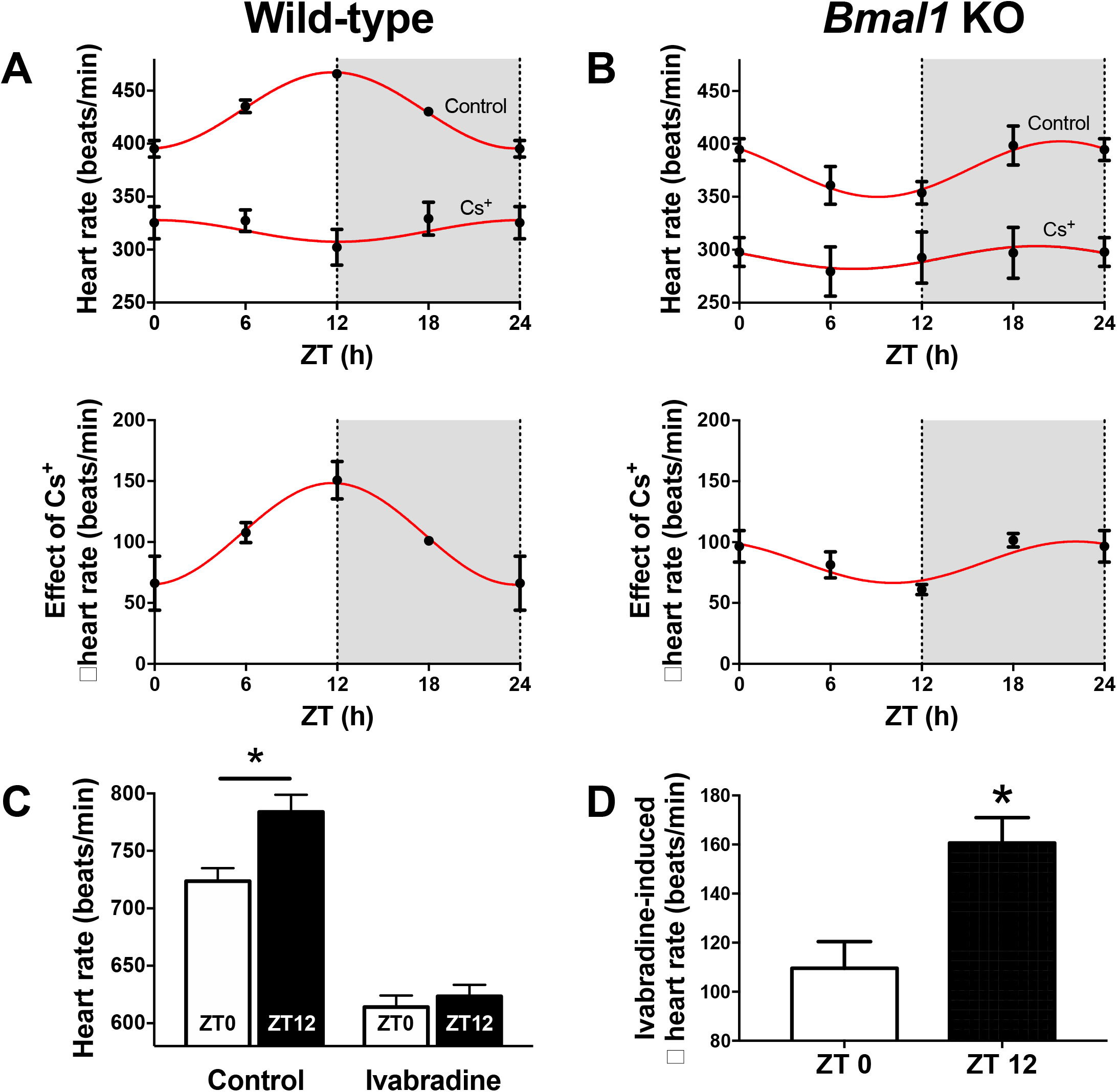
Effect of block of HCN4 and *I*_f_ on intrinsic heart rate and heart rate *in vivo*. **A**, Intrinsic heart rate before and after the application of 2 mM Cs^+^ (top) and change in intrinsic heart rate after application of Cs^+^ (bottom) measured in the isolated sinus node from wild-type mice (n=10 mice for ZT 0; n=5 for ZT 6; n=9 for ZT 12; n=4 for ZT 18). **B**, Intrinsic heart rate before and after the application of 2 mM Cs^+^ (top) and change in intrinsic heart rate after application of Cs ^+^ (bottom) measured in the isolated sinus node from cardiac-specific *Bmal1* knockout mice (n=8 mice for ZT 0; n=5 for ZT 6; n=4 for ZT 12; n=3 for ZT 18). **C**, *In vivo* heart rate of wild-type mice measured at ZT 0 and ZT 12 before and after the administration of 6 mg/kg ivabradine (n=5 mice for ZT 0; n=5 for ZT 12). *P<0.05; two-way ANOVA with Dunnett’s post-hoc test. **D**, Change in *in vivo* heart rate after administration of ivabradine (from same experiments as for C). *P<0.05; two-tailed paired t-test.

The so-called membrane and Ca^2+^ clocks are known to be responsible for pacemaking in the sinus node^22^. The membrane clock comprises ionic currents carried by a variety of ion channels. Although *I*_f_ carried by HCN channels is considered to be the most important, other ion channels also play a significant role. Various K^+^ channels showed a circadian rhythm (Figure 2G) and their potential role is considered in the Discussion. The Ca^2+^ clock involves spontaneous Ca^2+^ releases (Ca^2+^ sparks) from the sarcoplasmic reticulum during diastole. This drives Ca^2+^ extrusion via the electrogenic Na^+^-Ca^2+^ exchanger (*Slc8a1*; NCX1). The resulting inward current generated by NCX1 helps drive pacemaking. Although a circadian rhythm was not detected in any of the principal components of the Ca^2+^ clock (apart from Ca^2+^/calmodulin-dependent protein kinase IIδ – *Camk2d*), the potential role of the Ca^2+^ clock in the circadian rhythm in the intrinsic heart rate was investigated by recording Ca^2+^ sparks from isolated sinus node cells and measuring the effect of incapacitating the Ca^2+^ clock by 2 μM ryanodine on the intrinsic heart rate measured in the isolated sinus node (Figures S4-S6). Measurement of Ca^2+^ sparks showed that at ZT 12 as compared to ZT 0 there was an increase in spark duration and a non-significant increase in spark amplitude and consequently there was an increase in spark mass (calculated as amplitude×1.206×spark full width at half maximum amplitude^3^)^23^ (P=0.056), although there was a decrease in spark frequency and diastolic Ca^2+^ (Figure S5). The increase in spark mass at ZT 12 could increase the impact of the Ca^2+^ clock on pacemaking (although the effect could be mitigated by the reduction of Ca^2+^ spark frequency). In the isolated sinus node, the effect on the intrinsic heart rate of incapacitating the Ca^2+^ clock by ryanodine was greater at ZT 12 than ZT 0 (Figure S6A). In the presence of ryanodine, the circadian rhythm in the intrinsic heart rate was abolished (Figure S6A, top) and it is concluded that the Ca^2+^ clock also participates in the circadian rhythm in the intrinsic heart rate.

## Link between local circadian clock in sinus node and membrane and Ca^2+^ clocks

To investigate a possible link between the local circadian clock in the sinus node and the circadian rhythm in the intrinsic heart rate, experiments were conducted on a transgenic mouse in which the *Bmal1* gene had been knocked out in the heart only (driven by the α myosin heavy chain promoter)^24^. Knockout of *Bmal1* is known to incapacitate the circadian clock^25^. Figure 6A confirms that *Bmal1* mRNA was effectively knocked out in the sinus node at both ZT 0 and ZT 12 and Figure 6B shows that this disrupted the circadian rhythm in the expression of *Clock* mRNA – evidence that the circadian clock in the sinus node had been disrupted as expected. In the cardiac-specific *Bmal1* knockout mouse, from ZT 0 to ZT 12, there was no longer a significant variation in expression of *Hcn4* mRNA (Figure 6C) and HCN4 protein (Figure 3D,E). Consistent with this, the variation in *I*_f_ density from ZT 0 to ZT 12 was blunted (Figure 6D,E); this is best shown by Figure 6F, which compares the variation in *I*_f_ density from ZT 0 to ZT 12 in wild-type and cardiac-specific *Bmal1* knockout mice. Finally, Figure 5B (top) shows the intrinsic heart rate as measured in the isolated sinus node: whereas there was a circadian rhythm in the wild-type mice with the intrinsic heart rate peaking at ~ZT 12 (Figure 5A, top), in cardiac-specific *Bmal1* knockout mice this normal pattern was lost (Figure 5B, top). Furthermore, whereas there was a circadian rhythm in the reduction of the intrinsic heart rate on block of *I*_f_ by Cs^+^ in wild-type mice (with the effect peaking at ~ZT 12; Figure 5A, bottom), in cardiac-specific *Bmal1* knockout mice, once again, this pattern was lost (Figure 5B, bottom). In contrast, cardiac-specific *Bmal1* knockout had little effect on the circadian rhythm in the Ca^2+^ clock-control of the intrinsic heart rate (Figure S6B). It is concluded that the local circadian clock in the sinus node is controlling HCN4 and *I*_f_ (but perhaps not the Ca^2+^ clock) and thereby the intrinsic heart rate.

**Figure 6.**
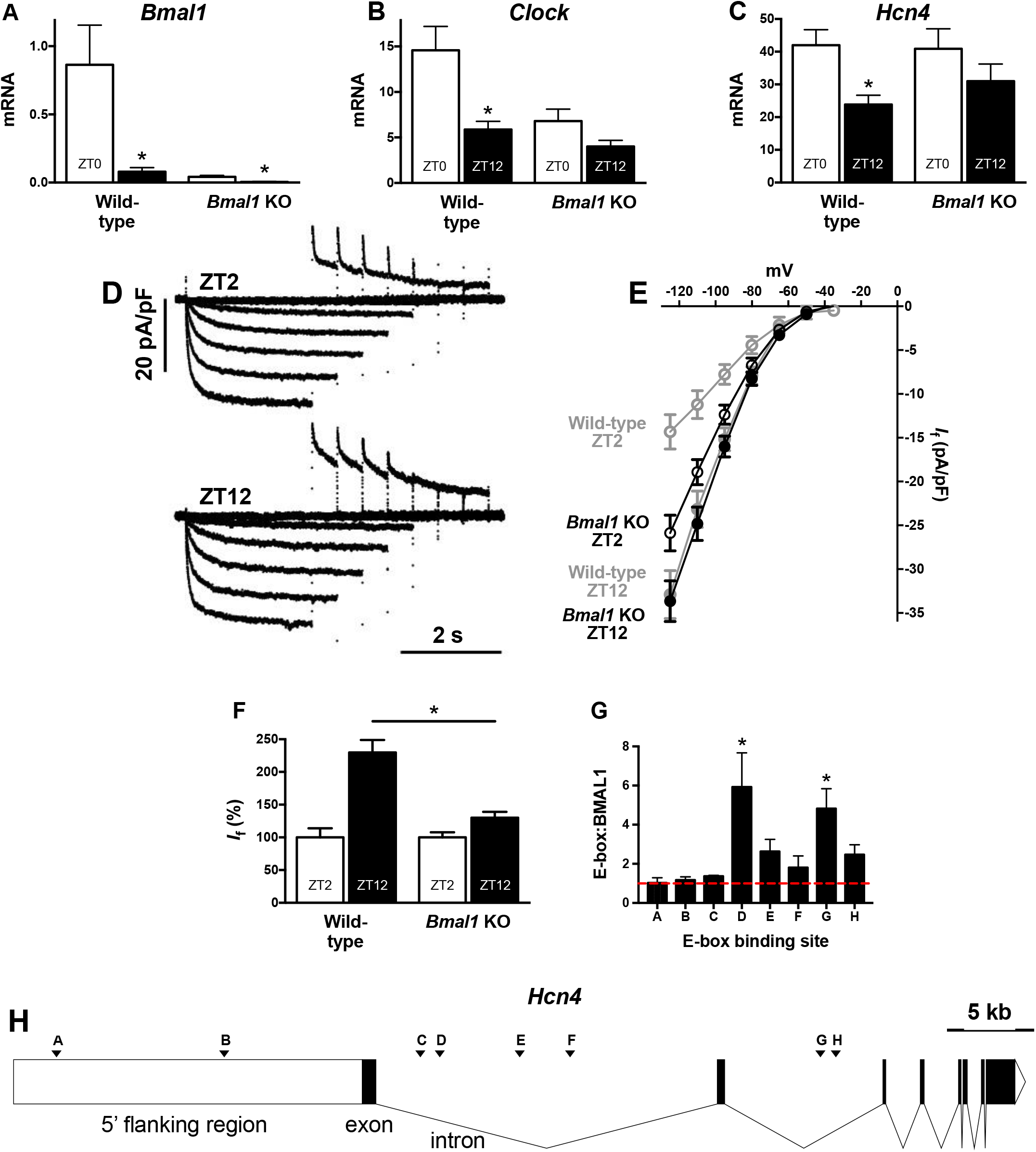
Cardiac-specific knockout of *Bmal1* stops or blunts circadian rhythm in *Hcn4* and *I*_f_. **A**, **B** and **C**, Expression of *Bmal1*, *Clock* and *Hcn4* mRNA in the sinus node at ZT 0 and ZT 12 in the sinus node of wild-type mice (n=7 mice for ZT 0 and ZT 12) and cardiac-specific *Bmal1* knockout mice (n=7 mice for ZT 0 and ZT 12). *ZT 12 versus corresponding ZT 0, P<0.05; two-way ANOVA with Tukey’s post-hoc test. **D**, Families of recordings of *I*_f_ made at ZT 2 and ZT 12 from sinus node cells isolated from cardiac-specific *Bmal1* knockout mice. **E**, Current-voltage relationships for *I*_f_ recorded at ZT 2 and ZT 12. Data shown in black were obtained from cardiac-specific *Bmal1* knockout mice (n=15 cells/3 mice at ZT 0; n=16 cells/3 mice at ZT 2). Data shown in grey were obtained from wild-type mice and have already been shown in Figure 4B. **F**, Density of *I*_f_ recorded at ZT 2 and ZT 12 from sinus node cells isolated from wild-type mice (n=16 cells/4 mice for ZT 2; n=16 cells/5 mice for ZT 12) and cardiac-specific *Bmal1* knockout mice (n=14 cells/3 mice for ZT 2; n=15 cells/3 mice for ZT 12). *I*_f_ expressed as a percentage of *I*_f_ at ZT 0. *P<0.05; two-way ANOVA with Holm-Šídák post-hoc test. **G**, Eight potential E-box binding sites pulled down on immunoprecipitation of His-tagged BMAL1 from cells transfected with His-tagged *Bmal1*. Data are normalised to immunoprecipitation from untransfected control cells; the red dashed line equals one and is the baseline level. *binding site of interest versus binding site A, P<0.05; n=2; one-way ANOVA with Dunnett’s post-hoc test. **H**, Diagram of the *Hcn4* gene and 20 kb of the 5′ flanking region showing the position of the eight potential E-box binding sites, A-H.

## Evidence that clock protein, BMAL1, controls *Hcn4* transcription

The CLOCK:BMAL1 heterodimer acts as a transcriptional activator or enhancer by binding to E-box binding sites in the promoter, intron or exon of a gene^26,27^. Using RVISTA (https://rvista.dcode.org) we found eight canonical E-box binding sites in the *Hcn4* gene and 20 kb of its 5′ flanking region (Figure 6H). *In vitro* ChIP was used to test whether BMAL1 specifically binds to these sites: 3T3-L1 cells were UV cross-linked following transfection with His-tagged *Bmal1*. His-tagged BMAL1 bound to its DNA targets was then immunoprecipitated using antibodies directed against the His-tag. DNA pulled down by ChIP was analysed by qPCR using primers mapping to the various E-box binding sites (Figure 6G). Figure 6G shows E-box binding sites D and G (within introns of the *Hcn4* gene; Figure 6H) were significantly more abundant than background signal (red dashed line).

## Role for the autonomic nervous system in circadian rhythm in heart rate i*n vivo*?

This study highlights the role of intrinsic factors in the circadian rhythm in intrinsic heart rate. Nevertheless, a role for the autonomic nervous system must be considered. In this study, *in vivo*, the heart rate was high during the awake period when the mice were physically active (Figure 1). If the level of physical activity is high, it would be expected to influence the heart rate via the autonomic nervous system (via an increase in sympathetic activity and possibly a decrease in vagal activity). However, in caged-housed mice with no access to a running wheel, activity levels will be low. A light pulse when mice are active is known to cause the mice to freeze and be inactive^12^. Figure 7A shows that a 1 h light pulse from ZT13 to ZT14 caused the physical activity of the mice to fall to baseline values, whereas the heart rate remained relatively high. In contrast, a 1 h light pulse from ZT1 to ZT2 was again associated with a baseline level of physical activity, but the heart rate was relatively low. Therefore, in this experiment, the heart rate was primarily influenced by the time of day rather than the activity level. Figure S7 shows that there is no discernible relationship between heart rate and physical level over 24 h (Figure 7A) and before, during and after the light pulses (Figure 7B). *In vivo*, the heart rate in the absence of physical activity can be obtained by recording the ECG from the anaesthetised mouse (although anaesthetic is known to depress the intrinsic pacemaker activity of the sinus node^28^ as well as cardiac vagal and sympathetic tone^29^). The heart rate in the anaesthetised mouse also varied in a daily manner – the heart rate was highest at ~ZT 12 and it varied by 51±15 beats/min over 24 h (Figure 7B; Table S1). It is concluded that in this study the effect of activity on the heart rate of the mice was slight and not discernible. Next, the effect of vagotomy was investigated. There are right and left vagal nerves and sectioning of both is lethal. However, vagal nerves to the sinus node are primarily, but not exclusively, from the right vagus^30^. The right vagus was sectioned in a group of rats (rat was studied, because we have experience of vagotomy in the rat^31^). Figure 7C shows that following the vagotomy the circadian rhythm in heart rate persisted.

**Figure 7.**
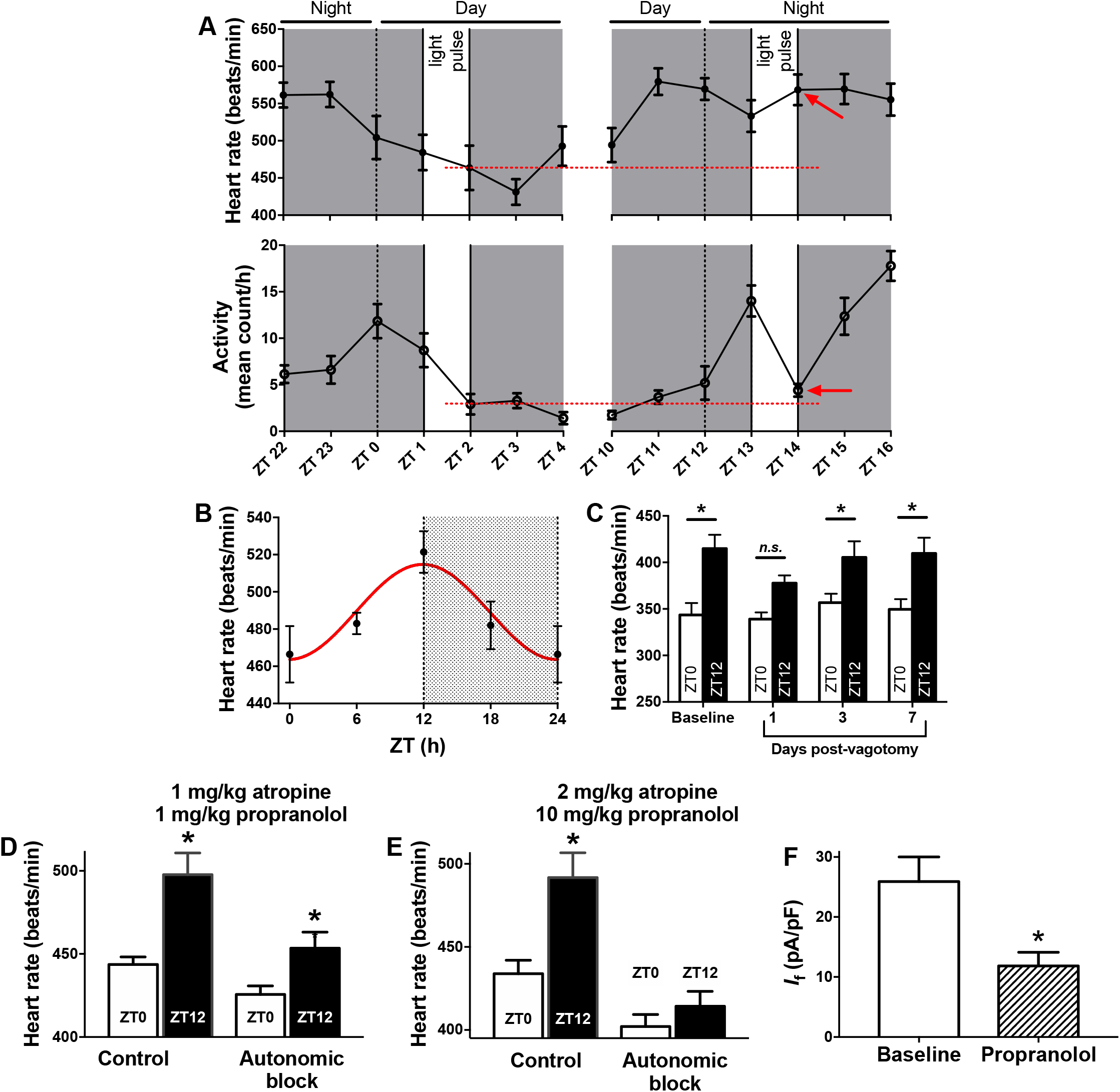
Circadian rhythm in heart rate *in vivo*. **A**, *In vivo* heart rate and physical activity measured by telemetry at the times shown during 24 h darkness (subjective day and night is shown) with the exception of a 1 h light pulse delivered towards the start of the day (left) or night (right). The dotted red lines highlight the heart rate and physical activity at the end of the day-time light pulse and the red arrows highlight the heart rate and physical activity at the end of the night-time light pulse. From the same experiment as Figure 1. **B**, *In vivo* heart rate measured from anaesthetised mice at four time points over 24 h (n=12 mice for ZT 0; n=10 for ZT 6; n=10 for ZT 12; n=10 for ZT 18). **C**, *In vivo* heart rate measured by telemetry at ZT 0 and ZT 12 in sham-operated and vagotomised rats at baseline (pre-surgery) and at 1, 3 and 7 days post-surgery *P<0.05. two-way ANOVA with Sidak’s multiple comparisons test. **D**, *In vivo* heart rate measured from anaesthetised mice at ZT 0 (n=9) and ZT 12 (n=9) before (Control) and after autonomic block by intraperitoneal injection of 1 mg/kg atropine and 1 mg/kg propranolol. *ZT 12 versus corresponding ZT 0, P<0.05; two-way ANOVA with Tukey’s post-hoc test. **E**, *In vivo* heart rate measured from anaesthetised mice at ZT 0 (n=5) and ZT 12 (n=5) before (Control) and after autonomic block by intraperitoneal injection of 2 mg/kg atropine and 10 mg/kg propranolol. *ZT 12 versus corresponding ZT 0, P<0.05; two-way ANOVA with Tukey’s post-hoc test. **F**, Density of *I*_f_ at −125 mV measured from 12 sinus node cells (from 3 mice) before and after application of 68.4 μM propranolol. *P<0.05; paired t test.

We further tested the involvement of the autonomic nervous system by blocking cardiac muscarinic and β receptors by atropine and propranolol. Experiments were conducted on anaesthetised mice, and intraperitoneal injections of 1 mg/kg atropine and 1 mg/kg propranolol were given, similar doses to those used by others^32–34^ and by us in an earlier study of exercise training-induced bradycardia in the mouse^35^. After autonomic blockade, the circadian rhythm in heart rate persisted, although it was reduced in amplitude (Figure 7D). In our earlier study, similar concentrations of atropine and propranolol also failed to abolish the exercise training-induced bradycardia^35^; we concluded that the autonomic nervous system is not responsible for the exercise training-induced bradycardia^35^ and based on Figure 7D we could conclude that it is also not solely responsible for the circadian rhythm in heart rate *in vivo*. However, following the publication of our study of exercise training-induced bradycardia, Aschar-Sobbi *et al.*^36^ used intraperitoneal injections of higher concentrations of atropine and propranolol (2 mg/kg atropine and 10 mg/kg propranolol) and stated that the exercise training-induced bradycardia in the mouse is abolished on autonomic blockade. Therefore, we repeated the experiment in Figure 7D, but we used intraperitoneal injections of 2 mg/kg atropine and 10 mg/kg propranolol as did Aschar-Sobbi *et al.*^36^. This time, after autonomic blockade, the circadian rhythm in heart rate was lost (Figure 7E). Although this suggests that the autonomic nervous system is involved in the circadian rhythm in heart rate *in vivo*, the effect of the higher concentrations of atropine and propranolol must be taken into consideration. Propranolol blocks β-receptors with an IC_50_ of 12 nM (http://www.selleckchem.com/products/propranolol-hcl.html). However, work on heterologously expressed ion channels in cell lines has shown that propranolol blocks the cardiac Na^+^ channel, Na_v_1.5 (*Scn5a*) with an IC_50_ of 2.7 μM^37^ and HCN4 with an IC_50_ of 50.5 μM^38^. It was confirmed that propranolol also blocks *I*_f_ in native heart cells: in mouse sinus node cells, 68.4 μM propranolol blocked *I*_f_ by 54% (Figure 7F); typical current traces and current-voltage relationships are shown in Figure S8. Assuming that only 10% of propranolol is free^39^, an injection of 10 mg/kg propranolol is estimated to give a propranolol concentration of 6 μM if it partitions in all body water (~60% of body mass in the mouse^40^) and 66 μM if it only partitions into the blood (58.5 ml/kg in the mouse; https://www.nc3rs.org.uk/mouse-decision-tree-blood-sampling). Therefore, based on these estimates, it is possible that 10 mg/kg propranolol may block, to some extent, Na_v_1.5 and HCN4, both of which vary in a circadian manner (Figure 2G). If propranolol did block HCN4 to a significant extent, it is not surprising that propranolol abolished the circadian rhythm in heart rate (Figure 7E), because ivabradine also did (Figure 5C). Propranolol at a dose of 1 mg/kg as used for Figure 7D is expected to cause less block of Na_v_1.5 and little or no block of HCN4 and yet complete block of β-receptors (the expected plasma concentration is still many times greater than the IC_50_ for block of β-receptors). As an aside, calculation shows that atropine at a dose of 1 or 2 mg/kg is sufficient for complete block of M2 muscarinic receptors. Based on Figure 7D, the autonomic nervous system cannot be solely responsible for the circadian rhythm in heart rate *in vivo*.

## DISCUSSION

For the first time, we show here that: (i) there is a circadian or daily rhythm in the *intrinsic* heart rate set by the pacemaker of the heart, the sinus node, (ii) there is a functioning circadian clock in the sinus node, (iii) there is a circadian rhythm in the expression of a variety of cardiac ion channels, including *Hcn4* and its corresponding ionic current, *I*_f_, in the sinus node, (iv) *Hcn4* transcription is directly controlled by the clock transcription factor, BMAL1, (v) the circadian rhythm in *Hcn4* plays an important role in the circadian rhythm in the *intrinsic* heart rate.

### Local circadian clock in heart

Previously, it has been demonstrated that there a functioning circadian clock in the heart^e.g.41^. This is the first report of a functioning circadian clock in the sinus node: qPCR showed the presence of transcripts for many key circadian clock components and many showed the expected circadian rhythm and expected phase relationship; for example, *Bmal1* was downregulated, but *Cry2*, *Per1* and *Per2* were upregulated at ZT 12 compared to ZT 0 (Figure 2C). We also showed a circadian rhythm in *Per1* using a bioluminescent reporter gene (Figure 2F).

### Circadian rhythm in cardiac ion channel expression is common

Here we show a circadian rhythm in *Hcn4*, Na_v_1.5 (*Scn5a*), K_v_1.4 (*Kcna4*), K_v_4.2 (*Kcnd2*), K_v_1.5 (*Kcna5*), ERG (K_v_11.1; *Kcnh2*), K_ir_6.1 (*Kcnj8*), K_ir_6.2 (*Kcnj11*), SUR1 (*Abcc8*), SUR2 (*Abcc9*), TASK-1 (K2p3.1; *Kcnk3*) and Cl^−^ voltage-gated channel 2 (*Clcn2*) in the sinus node (Figure 2G). Further experiments have shown that *Hcn1* also shows a circadian rhythm in the sinus node (Yanwen Wang, unpublished data). Curiously, *Hcn2* has been reported to show a circadian rhythm in liver^42^. A circadian rhythm in cardiac ion channel expression is not only seen in the sinus node – it is likely to be occurring throughout the heart. In atrial muscle, a circadian rhythm has been reported in K*v*4.2 (*Kcnd2*)^41,43,44^, KChIP2 (*Kcnip2*)^41,44^, K_v_1.5 (*Kcna5*)^41,43^, TASK-1 (K2p3.1; *Kcnk3*)^41^, Cx40 (*Gja5*)^45^ and Cx43 (*Gja1*)^45^. In ventricular muscle, a circadian rhythm has been reported in Na_v_1.5 (*Scn5a*)^46^, Ca_v_1.2 (*Cacna1c*)^47^, K_v_4.2 (*Kcnd2*)^41,43,48^, KChIP2 (*Kcnip2*)^41,44^, K_v_1.5 (*Kcna5*)^41,43^, ERG (K_v_11.1; *Kcnh2*)^48^, TASK-1 (K2p3.1; *Kcnk3*)^41^, Cx40 (*Gja5*)^45^ and Cx43 (*Gja1*)^45^. Some circadian varying ion channels are common to the sinus node and the atrial or ventricular muscle: Na_v_1.5 (*Scn5a*), K_v_4.2 (*Kcnd2*), K_v_1.5 (*Kcna5*), ERG (K_v_11.1; *Kcnh2*) and TASK-1 (K2p3.1; *Kcnk3*). Of these, all were significantly more highly expressed at ZT 12 than ZT 0 in the sinus node (Figure 2G); in atrial and ventricular muscle, there was a qualitatively similar circadian rhythm in Na_v_1.5 (*Scn5a*)^46^, K_v_4.2 (*Kcnd2*)^41,43,48^, K_v_1.5 (*Kcna5*; in ventricular muscle, but not atrial muscle)^41,43^ and ERG (K_v_11.1; *Kcnh2*)^48^. However, the opposite circadian rhythm in TASK-1 (K2p3.1; *Kcnk3*) has been reported in atrial and ventricular muscle^41^.

This study has demonstrated that *Hcn4* is likely to be under the control of BMAL1 generated by the local clock (Figure 6). In the ventricle, it has been reported that: Na*v*1.5 (*Scn5a*) and ERG (K_v_11.1; *Kcnh2*) are under the control of CLOCK:BMAL1 generated by the local clock^46,48^; and K_v_4.2 (*Kcnd2*), K_v_1.5 (*Kcna5*), TASK-1 (K2p3.1; *Kcnk3*), Cx40 (*Gja5*) and Cx43 (*Gja1*) are under the control of the suprachiasmatic nucleus (not the local clock) via the autonomic nervous system^41^. Confusingly, KChIP2 (*Kcnip2*) has been reported to be under the control of a BMAL1-dependent (i.e. local clock-dependent) oscillator, krüppel-like factor 15 (*Klf15*)^44^, and yet under the control of the suprachiasmatic nucleus (not the local clock) via the autonomic nervous system^41^.

### Circadian rhythm in intrinsic heart rate

Cardiac-specific knockout of *Bmal1* abolished the normal circadian rhythm in the intrinsic heart rate (Figure 5A,B), proving that it is under the control of the local circadian clock (however, it is interesting that some type of rhythm, albeit abnormal, remained after cardiac-specific knockout of *Bmal1* – Fig. 5B). We measured a circadian rhythm in HCN4 protein and *I*_f_ (Figures 3B-E, 4 and S2) roughly in phase with the circadian rhythm in the intrinsic heart rate (in contrast *Hcn4* mRNA was out of phase – Figure 3A; Table S1). It is likely that the circadian rhythm in *I*_f_ plays an important role in the intrinsic heart rate: computer modelling suggested that the changes in *I*_f_ are sufficient to explain the changes in intrinsic heart rate (Figure 4G); block of HCN4 and *I*_f_ by Cs^+^ abolished the circadian rhythm in the intrinsic heart rate (Figure 5A); and cardiac-specific knockout of *Bmal1* abolished the normal circadian rhythm in *Hcn4* mRNA and protein, *I*_f_ and the effect of Cs^+^, as well the intrinsic heart rate (Figures 3E, 5B and 6C,E,F). However, it is likely that *I*_f_ is not the only mechanism involved: there was a circadian rhythm in CaMKIIδ (Camk2d) and Ca^2+^ sparks (Figures 2G and S5), and incapacitating the Ca^2+^ clock with ryanodine eliminated the circadian rhythm in the intrinsic heart rate (Figure S6A). In addition, it is possible that the circadian rhythm detected in various K^+^ channels, particularly ERG (K_v_11.1; *Kcnh2*) (Figure 2G), also plays a role; in the sinus node, ERG (K_v_11.1; *Kcnh2*), responsible for the rapid delayed rectifier K^+^ current, *I*_K,r_, sets the maximum diastolic potential and blocking *I*_K,r_ abolishes pacemaking^49^.

### Circadian rhythm in heart rate *in vivo* – is it multifactorial?

This study has demonstrated a circadian rhythm in the *intrinsic* heart rate. Implications of this for the circadian rhythm in heart rate *in vivo* can be speculated on. Previously, the circadian rhythm in heart rate *in vivo* has been attributed to the autonomic nervous system and in particular to high vagal tone during the sleep period^4^. This primarily has been based on heart rate variability as a surrogate measure of autonomic tone^5^; however, as discussed above, this cannot be used in any simple way as a measure of autonomic tone^8^. We report that autonomic blockade utilising high concentrations of atropine and propranolol abolished the circadian rhythm in heart rate *in vivo* (Figure 7E). Abolition of the circadian rhythm in heart rate *in vivo* on autonomic blockade has also been reported by Tong *et al*.^41^ However, Oosting *et al.*^50^ have reported that autonomic blockade does not abolish the circadian rhythm in heart rate. Furthermore, caution must be used in interpreting the effect of autonomic blockade involving atropine and propranolol, because propranolol is non-specific and also blocks Na_v_1.5 (*Scn5a*) and *I*_Na_ and HCN4 and *I*_f_ (Figures 7F and S8)^37,38^. Because *I*_f_ at least is greater during the awake period, propranolol will be expected to have a greater depressing effect on heart rate during the awake period. This is an alternative explanation of why the circadian rhythm in heart rate *in vivo* was abolished by autonomic blockade in this study when utilising high concentrations of atropine and propranolol (Figure 7E). Nevertheless, if it is assumed that the autonomic nervous system is involved, the right vagus nerve (primarily, but not exclusively, responsible for the innervation of the sinus node^30^) cannot be solely responsible, because sectioning the right vagus did not impact the circadian rhythm in heart rate *in vivo* in the rat (Figure 7C). This suggests that a circadian rhythm in sympathetic nerve activity or plasma catecholamine levels is more important (if it is assumed that the autonomic nervous system is involved). However, it has been reported that transgenic knockout of the M2 receptor or β1, β2 and β3 adrenoceptors has no effect on the circadian rhythm in heart rate *in vivo*^51^. In addition, cardiac transplant patients with autonomic denervation have a preserved nocturnal bradycardia 7-36 months after transplantation^52,53^. This work does not support a role for the autonomic nervous system in the circadian rhythm in heart rate *in vivo*.

This study has shown that there is a circadian rhythm in the intrinsic heart rate of the appropriate amplitude (72-107 beats/min) and phase (peaking at ~ZT 12; Figures 2A and 5A; Table S1) to be able to explain the circadian rhythm in heart rate *in vivo* (Figure 1). Furthermore, block of HCN4 and *I*_f_ by ivabradine had a bigger effect on heart rate during the awake period and abolished the circadian rhythm in heart rate *in vivo* in the mouse (Figure 5C). Data consistent with this have been obtained by Ptaszynski *et al.*^54^ and Grigoryan *et al.*^55^ who studied the effect of ivabradine on the heart rate of patients either with inappropriate sinus node tachycardia or ischaemic heart disease and heart failure; whereas ivabradine caused a large decrease in heart rate during the day, it caused little decrease at night. In other words, the effect of ivabradine on heart rate showed a circadian rhythm and was greater when the subjects were awake. This is consistent with the work on the mice (Figure 5C,D)^54^. However, it is known that cardiac-specific knockout of *Bmal1*^46^ or cardiac-specific expression of a dominant negative *Clock* mutant^56^ does not abolish the circadian rhythm in heart rate *in vivo* (although it does reduce it). Furthermore, we show here that cardiac-specific knockout of *Bmal1* abolishes the *normal* circadian rhythm in *Hcn4* (mRNA and protein), *I*_f_ and intrinsic heart rate (Figures 3E, 5 and 6C,E,F). Therefore, HCN4 and *I*_f_ cannot be the only mechanism controlling the circadian rhythm in heart rate *in vivo* – perhaps the reduction of the circadian rhythm in heart rate *in vivo*^46^ on cardiac-specific knockout of *Bmal1* represents the contribution of the circadian rhythm in the intrinsic heart rate to the circadian rhythm in heart rate *in vivo*. What is responsible for the circadian rhythm in heart rate *in vivo* that remains after cardiac-specific *Bmal1* knockout? This could be due to the autonomic nervous system. Consistent with this, Figure 7D shows that the circadian rhythm in heart rate *in vivo* was reduced in amplitude after autonomic blockade (achieved with the lower doses of atropine and propranolol). However, the contribution may not be in the way originally conceived (acute control of heart rate via changes in ionic conductance): Tong *et al.*^41,45^ have shown that the circadian rhythm of K_v_4.2 (*Kcnd2*), KChIP2 (*Kcnip2*), K_v_1.5 (*Kcna5*), TASK-1 (K2p3.1; *Kcnk3*), Cx40 (*Gja5*) and Cx43 (*Gja1*) in the atria and ventricles is lost after autonomic blockade suggesting transcriptional regulation mediated by the autonomic nervous system. It is interesting that β-agonists affect both *Bmal1* and *Per2* in heart^57,58^.

### Summary

In summary, this study has shown that there is a circadian rhythm in the *intrinsic* heart rate and this is likely to contribute to the circadian rhythm in heart rate *in vivo*. Our findings provide new mechanistic insight into the fundamental question of why the heart rate of a mammal is lower when asleep and also explains the nocturnal occurrence of bradyarrhythmias^1,3,59–63^.

## METHODS

Extended methods are given in the Supplementary Information.

### Animals

The telemetry study was approved by the Norwegian Council for Animal Research, in accordance with the Guide for the Care and Use of Laboratory Animals from the European Commission Directive 86/609/EEC. Ethical approval for vagotomy in rats was from University College London, in accordance with the UK Animals (Scientific Procedures) Act 1986. All remaining procedures were approved by the University of Manchester and were in accordance with the UK Animals (Scientific Procedures) Act 1986. Experiments were conducted on adult male mice: C57BL/6J mice obtained from Harlan Laboratories; *Per1*∷LUC mice (in which luciferase expression is driven by the mouse *Per1* promoter and 5’-UTR elements^64^) on a *Cry1*^+/+^*Cry2*^+/+^ or *Cry1*^−/−^*Cry2*^−/−^ background; and cardiac-specific *Bmal1* knockout mice.

### Electrophysiology

ECGs or ECG-like electrograms were recorded for measurement of heart rate: from conscious unrestrained mice using either subcutaneously implanted radio telemetry transmitters or an ECGenie; from isofluorane-anaesthetised mice using a conventional three-lead system; from isolated Langendorff-perfused hearts; and from isolated right atrial preparations encompassing the sinus node. In the Langendorff-perfused hearts, a suitable pacing protocol was also used to determine the corrected sinus node recovery time. Strips of sinus node tissue were dissociated into single cells by a standard enzymatic and mechanical procedure and *I*_f_ was investigated using the patch clamp technique in the whole-cell mode. A previously developed biophysically-detailed mathematical model of the mouse sinus node cell action potential^65^ was used to assess the effect of changes in the density of *I*_f_ on the spontaneous action potential.

### mRNA and protein expression

RNA was isolated from frozen 1 mm sinus node punch biopsies and gene expression measured using quantitative PCR (qPCR) either using medium throughput custom-designed Taqman Low Density Array Cards or individual SYBR green assays. HCN4 protein was investigated using either sinus node homogenates and western blotting or tissue cryosections and immunohistochemistry. *Per1* expression was also measured using the *Per1*∷LUC mouse: total bioluminescence was recorded for 96 h from freshly dissected intact sinus node preparations from *Per1*∷LUC mice using a photomultiplier tube assembly.

### *In vitro* chromatin immunoprecipitation (ChiP)

ChIP was performed on 3T3-L1 cells using the SimpleChIP chromatin immunoprecipitation kit according to the manufacturer’s instructions. Prior to cross-linking, 3T3-L1 cultures at a density of 10^4^ cells/cm^2^ were transfected with plasmid containing His-tagged *Bmal1* for 48 h. Antibody directed against the His-tag was used for immunoprecipitation. DNA obtained from ChIP was analysed by qPCR using primers mapping to canonical E-box binding sites, i.e., consensus binding sites for the CLOCK∷BMAL1 heterodimer. E-box binding sites on the HCN4 gene and in 20 kb of the 5′ flanking region were obtained using the RVISTA function within ECR browser (http://ecrbrowser.dcode.org/). Quantification of *Hcn4* E-box binding sites was conducted by, first, normalisation to the housekeeping genes *Gapdh* and *L7* and, secondly, normalisation to data from 3T3-L1 cells subjected to transfection treatment without plasmid

## ACKNOWLEDGEMENTS

We thank Ms Rayna Samuels for technical assistance.

## AUTHOR CONTRIBUTIONS

Mark R. Boyett, Alicia D’Souza and George Hart conceived the project. Mark R. Boyett, Alicia D’Souza, Hugh D. Piggins, Halina Dobrzynski and Elizabeth Cartwright obtained funding for the project. Anne Berit Johnsen, Alicia D’Souza, Ulrik Wisloff, Eleanor Gill, Charlotte Cox, Yanwen Wang and Nick Ashton were involved in biotelemetry recordings. Yanwen Wang carried out electrophysiological experiments with assistance from Dario DiFrancesco and Annalisa Bucchi in patch clamp recordings. Alicia D’Souza carried out transcriptional profiling, bioluminescence recording and autonomic block experiments with assistance from Cali Anderson, Sven Wegner and Sunil Logantha. Alicia D’Souza, Yu Zhang and Yanwen Wang carried out western blotting experiments. Yanwen Wang and Nicholas Black performed immunofluorescent labelling studies. Haibo Ni and Henggui Zhang performed computer modelling studies using experimental data collected by Yanwen Wang. Yanwen Wang was responsible for the generation, breeding and maintenance of transgenic animals and all experiments on these animals (with assistance from Cheryl Petit, Hugh D. Piggins and Alicia D’Souza). Serve Olieslagers, Alicia D’Souza and Paula da Costa Martins were responsible for chromatin immunoprecipitation experiments. Svetlana Mastitskaya performed vagotomy experiments. Mark R. Boyett and Alicia D’Souza produced the first version of the manuscript, although subsequently all authors contributed.

