## Supplement for "Circadian control of intrinsic heart rate via a sinus node clock and the pacemaker channel"

### Supplementary Methods

#### Experimental animals

In Norway, the telemetry study was approved by the Norwegian Council for Animal Research, in accordance with the Guide for the Care and Use of Laboratory Animals from the European Commission Directive 86/609/EEC. In the UK, care and use of laboratory animals conformed to the UK Animals (Scientific Procedures) Act 1986. Ethical approval for vagotomy in rats was granted by University College London and ethical approval for all other experimental procedures was granted by the University of Manchester. Mice were housed five per cage in a temperature-controlled room (22°C) with a 12 h:12 h light:dark lighting regime (unless stated otherwise) and free access to food and water. Lights-on was at zeitgeber time (ZT) 0 and lights-off at ZT 12. Mice were adapted to a lighting regime for at least two weeks before use. Mice were sacrificed by cervical dislocation at defined time points, e.g. ZT 0; during the dark period mice were handled with the aid of night vision goggles to prevent exposure of the mice to light. The majority of experiments were carried out using male 10-12 week old (initial age) 25-30 g C57BL/6J mice from Harlan Laboratories. *Cry*$^{+/-}$*Cry2*$^{+/-}$ mice[1] that had been bred with transgenic knock-in animals (*Per1*::LUC, in which luciferase expression is driven by the mouse *Per1* promoter and 5'-UTR elements[2]) were originally obtained from Professor G.T. van der Horst (Erasmus MC, Rotterdam, Netherlands). Adult male offspring mice lacking the core molecular circadian clock (deficient in the *Cryptochrome* genes *Cry1*$^{-/-}$*Cry2*$^{-/-}$) and their congenic male wild-type (*Cry1*$^{+/+}$*Cry2*$^{+/+}$) littermates in which the molecular clock is fully functional were used at 4-6 months old as described previously[3]. Finally, male 10-14 week old cardiac-specific *Bmal1* knockout mice were used. They were generated by breeding *Bmal1*$^{flox/flox}$ mice with α-myosin heavy chain-Cre mice. *Bmal1*$^{flox/flox}$ mice have loxP sites flanking exon 8 of the *Bmal1* gene (Jackson Labs 007668)[4]. In α-myosin heavy chain Cre mice (from Dr. Elizabeth Cartwright's laboratory),[5] the cardiac-specific α-myosin-heavy chain (*Myh6*) promoter drives expression of Cre recombinase. In the resulting offspring, the exon encoding the *Bmal1* basic helix-loop-helix (bHLH) domain was deleted in the Cre recombinase-expressing cardiac tissues. Details of rats used are given below.

#### ECG and activity recording via telemetry in mice

Mice were anaesthetised with isoflurane (2.5% in $O_2$) and sterilised remote radio telemetry transmitters (model TA11ETA-F10, Data Sciences International, Netherlands) were implanted into the peritoneal cavity. Two electrodes were subcutaneously placed on the left caudal rib region and right pectoral muscle. Following surgery, mice were single-housed at 37°C for 2 h and then transferred to the normal holding area for one week to allow recovery. The ECG and activity were continuously recorded for six days. Animals were maintained in a normal 12 h light/12 h dark cycle for the first three days and in constant darkness for the remainder of the experiment, during which 1 h light pulses were given at ZT 1-2 and ZT 13-14 on Day 5. Telemetry data were received by small animal receiver RPC-1 (Data Sciences International), amplified by a Data Exchange Matrix (Data Sciences International) and acquired by Data Quest A.R.T. (P3 Plus version 4.8, Data Sciences International)[6]. The ECG was analysed using LabChart 7 (ADInstruments). ECG data shown in Fig. 1A are smoothed (2 h moving average). For activity data, 12 minutes every hour were analysed and mean activity was averaged per hour across animals. Activity readings (distance each mouse travelled) were taken every 5 min. For each mouse, for each hour, the mean of 12 such readings was recorded.

#### Biotelemetry and vagotomy in rats

Biotelemetry was used to record systemic arterial blood pressure and heart rate in vagotomised male 300-350 g Sprague-Dawley rats. Animals were anaesthetised with isoflurane (3% in $O_2$) and a laparotomy was performed to expose the abdominal aorta. A catheter connected to a telemetry pressure transducer (model TA11PAC40, DSI) was advanced rostrally into the aorta and secured with Vetbond (3M). The transmitter was secured to the abdominal wall and the incision was closed by successive suturing of the abdominal muscle and skin layers. Carprofen was given and the animals were returned to their home cages where they were allowed to recover for at least seven days. After the recovery period of one week and following recordings of the baseline haemodynamic parameters for 24 h, the animals were anaesthetised with isoflurane (3% in $O_2$). Using an aseptic technique, an anterior midline neck incision was performed, the sternohyoid and sternocleidomastoid muscles were retracted and the right cervical vagus nerve was isolated and sectioned. Sham-operated rats underwent the same surgical procedures to expose the nerve, but the vagus was left intact. Heart rate was recorded for seven days after surgery.

**ECG recording from the anaesthetised mouse**

Mice were anaesthetised with isoflurane (2% in $O_2$). A three-lead ECG was recorded using LabChart 7. Positive and negative electrodes were placed subcutaneously on the right fore limb and hind limb, respectively, and a reference electrode was placed subcutaneously on the left hind limb. ECG parameters were measured using LabChart 7 and GraphPad Prism. After 10 min (to allow the heart rate to stabilise), the ECG was recorded for 2 min and the average heart rate calculated. Block of the autonomic nervous system was achieved by intraperitoneal injection of propranolol (1 mg /kg or 10 mg/kg) followed by intraperitoneal injection of atropine (1 mg /kg or 2 mg/kg) after ~15 min or when the effect of sympathetic block with propranolol had reached steady-state. After ~5 min (once a steady-state post atropine administration had been reached), the ECG was recorded for a further 5 min and the average heart rate calculated.

**Electrogram recording from the Langendorff-perfused heart**

Following sacrifice, the heart was removed and the aorta cannulated and retrogradely perfused with Tyrode's solution (containing in mM: 120 NaCl; 4 KCl; 1.3 $MgSO_4$; 1.2 $NaH_2PO_4$; 25.2 $NaHCO_3$; 5.8 glucose; 1.8 $CaCl_2$; equilibrated with 95% $O_2$ and 5% $CO_2$ to give a pH of 7.4) at 36±0.5°C at a flow rate 3.5-4 ml/min. An ECG-like electrogram was recorded by placing two electrodes on the left atrium and ventricular apex. Recordings were amplified using a Neurolog system (Digitimer) with low-pass and high-pass filters adjusted to optimise the signal-to-noise ratio. After cannulation, hearts were allowed to stabilise for 15 min and then the electrogram was recorded over 100 consecutive beats. The function of sinus node was evaluated by the corrected sinus node recovery time (cSNRT)[7]. First, the spontaneous cycle length (SCL; time between spontaneous heart beats) was measured. Next the heart was paced at three cycle lengths (SCL-10 ms, SCL-20 ms and SCL-30 ms) for 30 s at each cycle length by programmed electric pacing using Spike2 software. The duration between the last paced beat and the first spontaneous beat was measured as the sinus node recovery time (SNRT) and the cSNRT was calculated as SNRT minus SCL. The electrogram was recorded and analysed using LabChart 7 and GraphPad Prism.

**Electrogram recording from the isolated sinus node**

Following sacrifice, the heart was removed and the rear wall of the right atrium encompassing the sinus node was rapidly dissected. The preparation was superfused with Tyrode's solution (containing in mM: 100 NaCl, 4 KCl, 1.2 $MgSO_4$, 1.2 $KH_2PO_4$, 25 $NaHCO_3$, 1.8 $CaCl_2$; 10 glucose; equilibrated with 95% $O_2$ and 5% $CO_2$ to give a pH of 7.4) at 37°C at a flow rate of 10 ml /min. An electrogram was recorded using three electrodes: two electrodes pinned either side of the preparation and a third earth electrode. Recordings were amplified using the Neurolog system with low-pass and high-pass filters adjusted to optimise the signal-to-noise ratio. Electrograms were recorded and analysed using LabChart 7 and GraphPad Prism. After dissection, the preparation was allowed to stabilise for 15 min, after which the electrogram was recorded and the average beating rate calculated over 10 min. The superfusing solution was changed to Tyrode's solution containing 2 mM CsCl, 2 µM ryanodine or 10 µM (±)-isoproteronol. After 10 min of treatment, the electrogram was again recorded and the average beating rate calculated over 10 min when the beating rate was steady.

**RNA isolation and qPCR**

A ~1 mm biopsy was collected from the sinus node at the level of the main branch of the crista terminalis, rapidly frozen in liquid $N_2$ and stored at −80°C until use. Total RNA was isolated using an RNeasy Micro kit (Qiagen) and reverse transcribed to produce cDNA using a SuperScript VILO cDNA Synthesis Kit (Applied Biosystems). RNA purity and quantity was determined using a NanoDrop ND-1000 spectrophotometer (NanoDrop Technologies, Wilmington, DE, USA). qPCR was carried out using an ABI Prism 7900 HT Sequence Detection System (Applied Biosystems/Life Technologies Corporation, Carlsbad, CA, USA). The expression level of 88 transcripts was measured using custom-designed Taqman Low Density Array Cards (Applied Biosystems, catalogue number 4342259; format 96A; transcripts studied are listed in Table S2) as previously described[8] and according to the manufacturer's instructions. Briefly, 150 ng of cDNA and TaqMan Universal Master Mix (Applied Biosystems) were loaded into each port of microfluidic cards containing fluorogenic TaqMan probes and primers. Thermal cycling conditions were 50°C for 2 min and 94.5°C for 10 min, followed by 40 cycles at 97°C for 30 s and 59.7°C for 1 min. Data were collected with ABI Prism 7900 HT Sequence Detection System software (SDS 2.3) and analysed using RealTime Statminer (v4.1, Integromics). Expression levels of the housekeeping genes, *18s*, *Gapdh*, *Hmbs*, *Ipo8*, *Pgk1* and *Tbp*, were tested. *Ipo8* and *Tbp* were selected as optimal (most stable) endogenous controls using geNorm, NormFinder and Minimum Variance Median via RealTime Statminer. Transcript expression levels were calculated using the ΔCt method. Statistical significance was tested using the non-parametric Limma test and Benjamini–Hochberg False discovery rate correction set at 5%. The FDR-adjusted P values are shown in Table S2. For single assays, the PCR reaction mixture comprised 1 µl cDNA, 1× Qiagen assay (*Bmal1*, QT00101647; *Clock*, QT00197547; *Hcn4*, QT00268660; *Ipo8*, QT00291977; *Tbp*, QT00198443), 1× SYBR Green Master Mix (Applied Biosystems) and DNase-free water. All samples were run in duplicate. The reaction conditions were: denaturation step of 95°C for 10 min followed by 40 cycles of amplification and quantification steps of 95°C for 30 s, 60°C for 30 s and 72°C for 1 min. The melt curve conditions were: 95°C for 15 s, 60°C for 15 s and 95°C for 15 s. mRNA expression normalised to the expression of the housekeepers, *Ipo8* and *Tbp*, was calculated using the ΔCt method.

**_Per1_::LUC bioluminescence experiments**
Total bioluminescence was recorded for 96 h from individual intact sinus node preparations using a photomultiplier tube (H8259/R7518P, Hamamatsu, Welwyn) using previously described methods[9]. 3.5 cm dishes were prepared under sterile conditions with a layer of autoclaved high-vacuum grease (Dow Corning Ltd., Coventry, UK) on the top rim, a Millicell insert (Millipore, Billerica, USA) and loaded with 1 ml recording medium consisting of DMEM (Invitrogen) supplemented with 10% foetal calf serum and 1% penicillin/streptomycin, 0.1 mM luciferin in autoclaved Milli-Q water. A freshly dissected beating sinus node was placed on the Millicell insert following which the dish was sealed with a glass coverslip and placed into the photomultiplier tube assembly housed in a light-tight incubator (GalaxyR+;RS Biotech, Irvine, UK) and maintained at 37°C. Photon counts were integrated for 599 s every 600 s. Bioluminescence data were detrended by subtracting a 24 h running average from the raw data and smoothed with a 3 h running average.

**Western blot**
Snap frozen sinus node biopsies were homogenised using MP FastPrep-24 and 2 ml tubes containing FastPrep metal bead lysing matrix (1.4 mm) in RIPA buffer (Sigma Aldrich). Bradford protein assay was used to estimate total protein concentration following which samples were denatured using 25% SDS-sample buffer (100 mM Tris-HCl, pH 6.8, 25% v/v glycerol, 10% v/v SDS, 10% v/v β-mercaptoethanol, 0.1% w/v bromophenol blue) and heating to 80°C for 10 min. Proteins were separated using a 7.5% stain-free SDS-polyacrylamide gel electrophoresis (PAGE) (Bio-Rad) system with PreSciccion Plus (Bio-Rad) protein standards running at 50 mV for ~50 min in SDS running buffer (25 mM Tris, 192 mM glycine, 0.1% SDS). Gels were imaged using ChemiDoc MP and proteins transferred to PVDF (polyvinyl difluoride) membranes using the Trans-Blot Turbo transfer system (Bio-Rad) at 15 V/0.3 mA for 15 min according to the manufacturer's instructions. Successful transfer was confirmed by imaging using the ChemiDoc MP. PVDF membranes were washed in Tris-buffered saline (TBS) and blocked in milk-TBS-Tween (5% w/v non-fat dried Marvel milk, TBS, 0.1% v/v Tween 20) and probed with the following primary antibodies for 1 h at room temperature with gentle rocking: rabbit polyclonal anti-HCN4 (Alomone Labs), 1:100; rabbit polyclonal anti-actin (Sigma Aldrich), 1:1000. After washing, membranes were

then probed with horseradish peroxidase (HRP)-linked secondary antibody (HRP-linked anti-rabbit IgG, Cell Signalling). Unbound secondary antibody was removed by washing in TBS-Tween following which membranes were treated with Clarity Western ECL substrate (Bio-Rad) and imaged. Samples from different time points were run on the same gel to ensure identical exposure conditions. The chemiluminescent signal intensity was normalised to the relative quantification of the corresponding intensity of actin. Data from replicates were averaged and normalised to signal at ZT 0.

**Immunohistochemistry**

Three control and three cardiac-specific *Bmal1* knockout mice were sacrificed at four time points (ZT 0, ZT 6, ZT 12 and ZT 18). Following sacrifice, the heart was removed and the rear wall of the right atrium encompassing the sinus node was rapidly dissected. The preparations were immersed in optimal cutting temperature compound, rapidly frozen in liquid $N_2$ and stored at -80$^{o}$C until use. The preparations were cryosectioned in a coronal plane to a thickness of 10 μm. Slides were stained using Masson's trichrome stain and the sinus node region was identified using the sinus node artery as a landmark. Two sinus node containing slides (four sections per slide) were chosen for each animal. Frozen sections were fixed in 10% formaldehyde, permeabilised with 0.1% Triton X-100 for 30 min, blocked with 1% bovine serum albumin and incubated with anti-HCN4 raised in rabbit (Alomone Labs; 1:200). Finally, sections were incubated with Cy3-conjugated secondary antibodies raised in donkey (Alomone Labs; 1:400). Images were collected on a Zeiss LSM5 PASCAL confocal microscope with a Plan Apochromat x40/1.0 oil objective lens. The confocal settings were as follows: pinhole, 1.02 airy unit; scan speed, 1 kHz unidirectional; format 1024 × 1024. Images were collected using the following settings: Cy3, 543 nm excitation and 640-690 nm emission. Four high power images were taken per section. HCN4 immunofluorescence was quantified using Volocity cellular imaging software and analysed by Image J. The sum pixel intensity for the HCN4-stained area was divided by the area to give a measurement of intensity per unit area.

**Sinus node cell isolation and patch-clamp**

Following sacrifice, the heart was removed and the sinus node rapidly dissected. Strips of nodal tissue were dissociated into single cells by a standard enzymatic and mechanical procedure. The enzyme solution contained collagenase IV (224 U/ml, Worthington), elastase (1.42 U/ml, Sigma-Aldrich) and protease (0.45 U/ml, Sigma-Aldrich)[10]. Isolated sinus node cells were transferred to $Ca^{2+}$-containing Tyrode's solution and stored at 4°C for the day of the experiment. $I_f$ was recorded using a patch electrode in whole-cell mode during superfusion of Tyrode's solution (containing in mM: 140 NaCl, 5.4 KCl, 1.8 $CaCl_2$, 1 $MgCl_2$, 5 HEPES-NaOH, 10 D-glucose, pH 7.4). $BaCl_2$ (1 mM) and $MnCl_2$ (2 mM) were added to avoid contamination from other ionic currents. The bath temperature was maintained at 35±0.5°C. The pipette solution contained (in mM): 130 K-aspartate, 10 NaCl, 2 $CaCl_2$ (pCa=7), 2 $MgCl_2$, 10 HEPES, 5 EGTA, 2 ATP($Na_2$), 0.1 GTP, 5 creatine phosphate, pH 7.2. $I_f$ was measured during potential steps to −35 to −125 mV from a holding potential of −35 mV and normalised to cell capacitance. Cell capacitance was calculated from the current measured in response to a -10 mV pulse from the holding potential. To test the effect of propranolol on $I_f$, 68.4 μM propranolol was applied to sinus node cells. $I_f$ was recorded from the same cell before and after propranolol application, and the data were normalised to cell capacitance. Data were acquired at 1 kHz using an Axopatch 200 amplifier and pClamp 8 (Molecular Devices, Sunnyvale, CA, USA). Data were analysed off-line using Clampfit 10 (Molecular Devices, Sunnyvale, CA, USA), Origin 8 (Origin Lab Corp., Northampton, MA, USA) and GraphPad Prism.

**$Ca^{2+}$ spark measurement**

Isolated sinus node cells were loaded with the membrane permeable acetoxymethyl (AM) ester form of the fluorescent $Ca^{2+}$ indicator, Fluo-8 (Cambridge Bioscience), at a final concentration of 2.5 µM (in Tyrode's solution) for 5 min at room temperature. The cells were then placed in a Petri dish with Tyrode's solution and kept at 4°C in the dark until use. The Petri dish with cells was placed onto a Nikon Eclipse Ti-S microscope and left for 5 min for acclimatisation before superfusion with Tyrode's solution at 37°C in the dark. $Ca^{2+}$ sparks were recorded with an Andor Revolution XD confocal system with a Yokogawa spinning disk confocal system. Laser excitation of Fluo-8 was at 488 nm and emitted light was collected using a 515 nm long pass filter. The images were captured at an acquisition rate of 6 ms for 3 min. The XY image stacks were acquired using Andor IQ2 software and were analysed using the automated $Ca^{2+}$ spark detection software, xySpark, and ImageJ. Data were processed by GraphPad Prism 7.

**Computational modelling**

A previously developed biophysically-detailed mathematical model of the mouse sinus node cell action potential[11] was used to assess the effect of changes in the density of $I_f$ on the spontaneous action potential. The densities of $I_f$ (at a potential of -140 mV) at different times over 24 h were normalised to the value at ZT 2. The ratios were subsequently used to scale the maximum conductance of $I_f$ in the mathematical model to simulate the time-dependent changes in density of $I_f$. For each simulation, the standard forward Euler integration scheme with a time step of 0.001 ms was used to solve the ordinary differential equation system and this was found to give stable solutions[11]. The initial conditions in all simulations were kept the same with the original model[11]. The cycle length (time between consecutive action potentials) was measured between the point of maximum upstroke velocity ($dV/dt_{max}$) of two consecutive spontaneous action potentials; from this the 'heart rate' was calculated. 20 s of activity was computed to ensure the model reached a steady-state (with differences in cycle length of two consecutive pairs of action potentials being <0.01 ms).

***In vitro* ChIP**

ChIP was performed on 3T3-L1 cells (ATCC, #CL-173) using the SimpleChIP chromatin immunoprecipitation kit (CST, #9005) according to the manufacturer's instructions. Prior to cross-linking, 3T3-L1 cultures at a density of $10^4$ cells/$cm^2$ were transfected with plasmid containing His-tagged *Bmal1* (pBMPC3; gift from Aziz Sancar, Addgene, plasmid #31367) for 48 h. Antibody directed against the His-tag (CST, #12698) was used for immunoprecipitation. DNA obtained from ChIP was analysed by qPCR using primers mapping to canonical E-box binding sites, i.e., consensus binding sites for the CLOCK::BMAL1 heterodimer. E-box binding sites on the *Hcn4* gene and in 20 kb of the 5′ flanking region were obtained using the RVISTA function within ECR browser (http://ecrbrowser.dcode.org/). Quantification of *Hcn4* E-box binding sites was conducted by, first, normalisation to the housekeeping genes *Gapdh* and *L7* and, secondly, normalisation to data from 3T3-L1 cells subjected to transfection treatment without plasmid. The following PCR primers were used:

| | |
|---|---|
| Forward primer for *Hcn4* E-box binding site A | CACCCTTGCCTTCCTTCTACTG |
| Reverse primer for *Hcn4* E-box binding site A | ACAAAGAAAGACTCTTAGTGTGCAGGT |
| Forward primer for *Hcn4* E-box binding site B | GGGCTCTACCACCTCCCAG |
| Reverse primer for *Hcn4* E-box binding site B | GTCTGTGGGCAGGTCCATG |
| Forward primer for *Hcn4* E-box binding site C | GGCCATCTCTTCTCTGTGTCTGT |
| Reverse primer for *Hcn4* E-box binding site C | TAAGTATGAACTTCAAACGGCCAA |
| Forward primer for *Hcn4* E-box binding site D | GGGCTCACAAAACATGGCA |
| Reverse primer for *Hcn4* E-box binding site D | CCCACCAGAGCCTAGACCC |
| Forward primer for *Hcn4* E-box binding site E | GGCCTGATAATTGTACTGGTGGT |
| Reverse primer for *Hcn4* E-box binding site E | GCATTAGCCAGCCCTGTTATG |
| Forward primer for *Hcn4* E-box binding site F | GCCAGGTACCTGGCTTCC |
| Reverse primer for *Hcn4* E-box binding site F | GAACAAAGACTCAAGTACCACTCCC |
| Forward primer for *Hcn4* E-box binding site G | TAGCCAGGGCCTGAGGC |
| Reverse primer for *Hcn4* E-box binding site G | GCATGGAATGCACGGGTAG |
| Forward primer for *Hcn4* E-box binding site H | CCTGAAGGTCCCGTTCTTGT |
| Reverse primer for *Hcn4* E-box binding site H | CAGTGACAGATTGGCTTGCC |

| | |
|---|---|
| Forward primer for *Gapdh* | CTTCCGTGTTCCTACCC |
| Reverse primer for *Gapdh* | ACCTGGTCCTCAGTGTAGCC |
| Forward primer for *L7* | GAAGCTCATCTATGAGAAGGC |
| Reverse primer for *L7* | AAGACGAAGGAGCTGCAGAAC |

### Limitations of the Study

This study used male mice or rats exclusively; clearly, it will be important to study females in the future. In the present study, the 'intrinsic heart rate' was measured using three different types of preparation. The intrinsic heart rate is the heart rate set by the sinus node in the absence of external influences and it should be the same regardless of what preparation is used to measure it, but this was not the case: the 'intrinsic heart rate' measured in anaesthetised mice after autonomic blockade was ~420 to 440 beats/min (Fig. 7D), whereas the 'intrinsic heart rate' measured in Langendorff-perfused hearts was ~270 to 380 beats/min (Fig. 2A) and the 'intrinsic heart rate' measured in the isolated sinus node was ~400 to 470 beats/min (Fig. 5A). The heart rate of the anaesthetised mouse after autonomic blockade was ~520 beats/min in a previous study from our laboratory[8]. The heart rates of the Langendorff-perfused mouse hearts are consistent with heart rates for this preparation reported in the literature (251 to 422 beats/min)[12-20]. The heart rates of the isolated mouse sinus node are consistent with previous measurements (300-500 beats/min) from our laboratory[8]. The reasons for the differences are not known. However, it is possible that the heart rate in anaesthetised mice after autonomic blockade is influenced by some unknown neurohumoral factor. In addition, anaesthetic is known to affect the electrical activity of the sinus node[21]. The different heart rates in the Langendorff-perfused heart (perfused by the coronary vessels) and isolated sinus node (superficially perfused) could be the result of some unknown factor released from the vessels of the Langendorff-perfused heart. In the present study, the intrinsic heart rate was measured in all three preparations to overcome the limitations of any one preparation.

**Table S1. Summary of circadian rhythms in heart rate, *Bmal1*, *Clock*, *Hcn4* and $I_f$.** The majority of the data were calculated from the fitting of a sine wave. The periodicity was assumed to be 24 h. [a]data based on a small number of time points over 24 h and, therefore, subject to greater error. [b]data based on means at ZT 0 and ZT 12 and, therefore, no time course data available. [c]data obtained by extrapolation and therefore, subject to error. [d]data obtained with isoflurane anaesthesia, which is known to depress sinus node pacemaking as well as autonomic tone, and, therefore, the changes in heart rate are likely to be depressed. Data are ranked according to the likely sequence of events.

**Table S1 continued.**

| Event | Source data | Circadian rhythm in heart rate (beats/min) | Time of peak (ZT) | Timeline |
|---|---|---|---|---|
| **Early events** | | | | |
| *Bmal1* mRNA[a] | Figure 2D | - | 0.5±1.1 | |
| *Clock* mRNA[a] | Figure 2E | - | 0.2±1.0 | |
| *Hcn4* mRNA[a] | Figure 3A | - | 20.0±0.4 | |
| **Later event** | | | | **Peripheral clock** |
| HCN4 protein – western blot[a] | Figure S2 | - | 8.3±1.6 | |
| HCN4 protein – immunohistochemistry[a] | Figure S2 | - | 11.0±1.3 | ***Hcn4*** |
| $I_f$ amplitude | Figure 4D | - | 10.9±0.7 | **HCN4** |
| $I_f$ density | Figure 4C | - | 10.0±0.3 | |
| **Late events – intrinsic heart rate (not subject to autonomic influence)** | | | | **Funny current** |
| Computed heart rate[c] | Figure 4G | 101±11 | 10.0±0.3 | |
| Isolated sinus node[b] | Figure 5A | 72±7 | 11.7±0.5 | |
| Langendorff-perfused heart[b] | Figure 2A | 107 | - | |
| **Late events – normal heart rate (subject to autonomic influence)** | | | | **Intrinsic heart rate** |
| Heart rate *in vivo* of anaesthetised mouse[ad] | Figure 7B<br>Figure 7D<br>Figure 7E | 51±15<br>22<br>58 | 11.9±1.4<br>-<br>- | |
| Heart rate *in vivo* of conscious mouse during 12 h light, 12 h dark regime | Fig. 1 | 76±4 | 13.1±0.2 | |
| **Late events – normal heart rate (subject to autonomic influence) but with altered lighting regime** | | | | **Normal heart rate** |
| Heart rate *in vivo* of conscious mouse during 24 h dark regime | Fig. 1 | 103±5 | 15.1±0.2 | |

**Table S2. Expression of mRNA for clock components, transcription factors, ion channels, $Na^+$-$K^+$ pump subunits, intracellular $Ca^{2+}$-handling molecules, gap junction channels and hypertrophy markers in the sinus node at ZT 0 and ZT 12.** Differences between ZT 0 and ZT 12 were tested using the limma test and the FDR-corrected P values are shown. Statistically significant differences (P<0.05) are highlighted in yellow. n=7/9.

| mRNA | Protein | ZT 0 (mean±SEM) | ZT 12 (mean±SEM) | FDR-adjusted P value |
|---|---|---|---|---|
| **Clock genes** | | | | |
| *Arid1A* | AT-rich interactive domain-containing protein 1A | 1.29±0.25 | 1.59±0.11 | 0.301 |
| *Bhlhe40* | Class E basic helix-loop-helix protein 40 | 0.63±0.18 | 1.75±0.34 | 0.031 |
| *Bhlhe41* | Class E basic helix-loop-helix protein 41 | 0.95±0.10 | 4.35±0.79 | 0.001 |
| *Bmal1* | Aryl hydrocarbon receptor nuclear translocator-like protein 1 | 0.91±0.15 | 0.07±0.01 | 1.92E-06 |
| *Bmal2* | Aryl hydrocarbon receptor nuclear translocator-like protein 2 | 0.02±0.003 | 0.05±0.02 | 0.977 |
| *Clock* | Circadian locomoter output cycles protein kaput | 5.17±1.49 | 2.15±0.28 | 0.099 |
| *Cry1* | Cryptochrome-1 | 0.51±0.05 | 0.40±0.04 | 0.298 |
| *Cry2* | Cryptochrome-2 | 0.40±0.04 | 1.25±0.25 | 0.002 |
| *Csnk1e* | Casein kinase I isoform epsilon | 0.99±0.10 | 1.54±0.25 | 0.307 |
| *Lhx1* | LIM/homeobox protein Lhx1 | 0.001±0.0003 | 0.001±0.0005 | 0.873 |
| *Per1* | Period circadian protein homolog 1 | 0.43±0.06 | 1.84±0.16 | 2.26E-05 |
| *Per2* | Period circadian protein homolog 2 | 0.18±0.02 | 0.93±0.05 | 5.11E-07 |
| *Per3* | Period circadian protein homolog 3 | 0.22±0.02 | 1.59±0.09 | 1.61E-08 |
| *Nr1d1* | Nuclear receptor subfamily 1 group D member 1 | 0.33±0.03 | 4.19±0.66 | 4.42E-08 |
| *Nr1d2* | Nuclear receptor subfamily 1 group D member 2 | 1.88±0.73 | 8.22±2.75 | 0.005 |
| *RoRa* | Nuclear receptor ROR-alpha | 1.57±0.17 | 2.19±0.27 | 0.186 |
| *Timeless* | Protein timeless homolog | 0.15±0.01 | 0.19±0.01 | 0.271 |
| **Circadian clock regulated transcription factors** | | | | |
| *Dbp* | D site-binding protein | 0.05±0.01 | 3.97±0.22 | 4.04E-11 |
| *Hlf* | Hepatic leukemia factor | 2.27±0.47 | 4.40±0.33 | 0.012 |
| *Tef* | Thyrotroph embryonic factor | 1.42±0.13 | 6.57±0.50 | 5.11E-07 |

**Table S2 continued.**

| Gene | Protein | ZT 0 (mean±SEM) | ZT 12 (mean±SEM) | FDR adjusted P value |
|---|---|---|---|---|
| | | **HCN channels** | | |
| *Hcn1* | HCN1 | 5.01±0.75 | 6.34±3.07 | 0.620 |
| *Hcn2* | HCN2 | 0.49±0.07 | 0.66±0.06 | 0.252 |
| *Hcn4* | HCN4 | 12.05±1.32 | 6.39±1.42 | 0.028 |
| | | **$Na^+$ channels** | | |
| *Scn1a* | $Na_v$1.1 | 0.002±0.0003 | 0.03±0.01 | 0.163 |
| *Scn5a* | $Na_v$1.5 | 4.17±0.45 | 8.08±0.83 | 0.005 |
| *Scn1b* | $Na_v$β.1 | 2.34±0.24 | 3.4±0.59 | 0.354 |
| | | **$Ca^{2+}$ channels** | | |
| *Cacna1c* | $Ca_v$1.2 | 1.71±0.27 | 2.08±0.35 | 0.928 |
| *Cacna1d* | $Ca_v$1.3 | 0.59±0.15 | 0.82±0.15 | 0.385 |
| *Cacna1g* | $Ca_v$3.1 | 1.12±0.14 | 1.17±0.13 | 0.789 |
| *Cacna1h* | $Ca_v$3.2 | 4.31±0.64 | 7.63±1.20 | 0.084 |
| *Cacna2d1* | $Ca_v$α2δ1 | 13.21±4.71 | 16.20±9.22 | 0.873 |
| *Cacna2d2* | $Ca_v$α2δ2 | 3.56±0.61 | 5±0.81 | 0.354 |
| *Cacnb2* | $Ca_v$β2 | 1.85±0.27 | 4.06±0.78 | 0.084 |
| | | **Transient outward $K^+$ channels** | | |
| *Kcna2* | $K_v$1.2 | 0.33±0.04 | 0.48±0.04 | 0.109 |
| *Kcna4* | $K_v$1.4 | 0.14±0.01 | 0.29±0.01 | 0.002 |
| *Kcnb1* | $K_v$2.1 | 1.70±0.18 | 1.23±0.11 | 0.163 |
| *Kcnd2* | $K_v$4.2 | 0.46±0.04 | 1.14±0.08 | 2.22E-04 |
| *Kcnd3* | $K_v$4.3 | 0.65±0.16 | 0.80±0.09 | 0.333 |
| *Kcnip2* | KChIP2 | 0.24±0.02 | 0.26±0.04 | 0.975 |
| | | **Delayed rectifier $K^+$ channels** | | |
| *Kcna5* | $K_v$1.5 | 0.40±0.05 | 0.73±0.06 | 0.021 |
| *Kcnh2* | ERG1 | 1.32±0.2 | 3.01±0.34 | 0.009 |
| *Kcnq1* | $K_V$LQT1 | 2.06±0.16 | 2.55±0.23 | 0.217 |
| | | **Inward rectifier $K^+$ channels** | | |
| *Kcnj2* | Kir2.1 | 1.55±0.15 | 1.78±0.18 | 0.596 |
| *Kcnj12* | Kir2.2 | 0.84±0.07 | 1.19±0.10 | 0.084 |
| *Kcnj4* | Kir2.3 | 0.004±0.0009 | 0.008±0.002 | 0.466 |
| *Kcnj14* | Kir2.4 | 0.11±0.02 | 0.10±0.01 | 0.629 |
| *Kcnj3* | Kir3.1 | 14.19±1.83 | 22.31±3.75 | 0.117 |
| *Kcnj5* | Kir3.4 | 7.60±0.65 | 10.06±0.95 | 0.187 |
| *Kcnj8* | Kir6.1 | 1.02±0.06 | 1.63±0.15 | 0.031 |
| *Kcnj11* | Kir6.2 | 1.32±0.14 | 2.86±0.26 | 0.002 |
| *Abcc8* | SUR1 | 1.57±0.19 | 2.88±0.27 | 0.006 |
| *Abcc9* | SUR2 | 2.23±0.33 | 4.67±0.26 | 0.002 |

**Table S2 continued.**

| mRNA | Protein | ZT 0 (mean±SEM) | ZT 12 (mean±SEM) | FDR adjusted P value |
|---|---|---|---|---|
| | | **Miscellaneous $K^+$ channels** | | |
| *Kcnn1* | SK1 | 0.16±0.02 | 0.29±0.07 | 0.157 |
| *Kcnn2* | SK2 | 0.48±0.05 | 0.60±0.04 | 0.217 |
| *Kcnn3* | SK3 | 0.08±0.01 | 0.10±0.01 | 0.328 |
| *Kcnk3* | TASK1 | 10.68±1.36 | 22.69±6.21 | 0.033 |
| *Trpc3* | TRPC3 | 1.85±0.04 | 0.12±0.01 | 0.489 |
| | | **$Cl^-$ channels** | | |
| *Cftr* | CFTR | 0.06±0.01 | 0.05±0.004 | 0.819 |
| *Clcn2* | CLC-2 | 0.08±0.01 | 0.30±0.04 | 0.001 |
| | | **$Na^+$-$K^+$ Pump** | | |
| *Atp1a1* | $Na^+$-$K^+$ pump α1 subunit | 45.33±9.48 | 42.64±8.96 | 0.983 |
| *Atp1a2* | $Na^+$-$K^+$ pump α2 subunit | 20.26±3.09 | 34.74±13.72 | 0.443 |
| *Atp1a3* | $Na^+$-$K^+$ pump α3 subunit | 0.2±0.05 | 0.62±0.14 | 0.106 |
| | | **Intracellular $Ca^{2+}$-handling molecules** | | |
| *Casq2* | Calsequestrin 2 | 59.30±22.84 | 40.26±16.71 | 0.385 |
| *Camk2d* | Camk2δ | 3.04±0.37 | 4.92±0.45 | 0.046 |
| *Slc8a1* | NCX1 | 9.30±0.49 | 15.28±2.76 | 0.179 |
| *Pln* | Phospholamban | 1.15±0.09 | 1.47±0.24 | 0.789 |
| *Ryr2* | RYR2 | 63.75±24.87 | 70.52±37.08 | 0.807 |
| *Ryr3* | RYR3 | 0.48±0.27 | 0.30±0.04 | 0.807 |
| *Sln* | Sarcolipin | 97.28±5.87 | 86.02±14.31 | 0.415 |
| *Atp2a2* | SERCA2a | 857±429.6 | 953.59±633.82 | 0.852 |
| | | **Gap junction channels** | | |
| *Gjd3* | Cx30.2 | 0.06±0.01 | 0.08±0.01 | 0.367 |
| *Gja5* | Cx40 | 1.15±0.22 | 2.29±0.52 | 0.102 |
| *Gja1* | Cx43 | 1.95±0.22 | 1.488±0.26 | 0.247 |
| *Gjc1* | Cx45 | 0.06±0.02 | 0.06±0.014 | 0.835 |

**Table S2 continued.**

| mRNA | Protein | ZT 0 (mean±SEM) | ZT 12 (mean±SEM) | FDR adjusted P value |
|---|---|---|---|---|
| | | **Transcription factors** | | |
| *Gata4* | GATA4 | 3.32±0.34 | 3.35±0.37 | 0.975 |
| *Hand2* | Heart- and neural crest derivatives-expressed protein 2 | 1.21±0.1 | 1.72±0.17 | 0.136 |
| *Isl1* | Insulin gene enhancer protein ISL-1 | 0.36±0.08 | 0.19±0.04 | 0.110 |
| *Klf4* | Krueppel-like factor 4 | 1.64±0.23 | 1.40±0.07 | 0.593 |
| *Klf15* | Krueppel-like factor 15 | 0.56±0.057 | 1.03±0.08 | 0.011 |
| *Mef2c* | Myocyte-specific enhancer factor 2C | 1.02±0.11 | 1.42±0.20 | 0.411 |
| *Nkx2.5* | Homeobox protein Nkx-2.5 | 1.36±0.18 | 1.60±0.31 | 0.693 |
| *Rest* | RE1-silencing transcription factor | 0.47±0.1 | 0.44±0.02 | 0.883 |
| *Pitx2* | Pituitary homeobox 2 | 0.14±0.03 | 0.12±0.01 | 0.620 |
| *Shox2* | Short stature homeobox protein 2 | 0.98±0.14 | 0.42±0.06 | 0.005 |
| *Srf* | Serum response factor | 2.46±0.15 | 2.30±0.15 | 0.695 |
| *Tbx3* | T-box transcription factor 3 | 1.80±0.22 | 0.96±0.12 | 0.028 |
| *Tbx5* | T-box transcription factor 5 | 8.57±0.8 | 11.49±1.51 | 0.376 |
| *Tbx18* | T-box transcription factor 18 | 0.78±0.14 | 0.41±0.06 | 0.063 |
| | | **Hypertrophy markers** | | |
| *Nppa* | Atrial natriuretic peptide | 114±22.17 | 317.57±103.9 | 0.032 |
| *Nppb* | Brain natriuretic peptide | 0.99±0.28 | 2.20±1.09 | 0.645 |

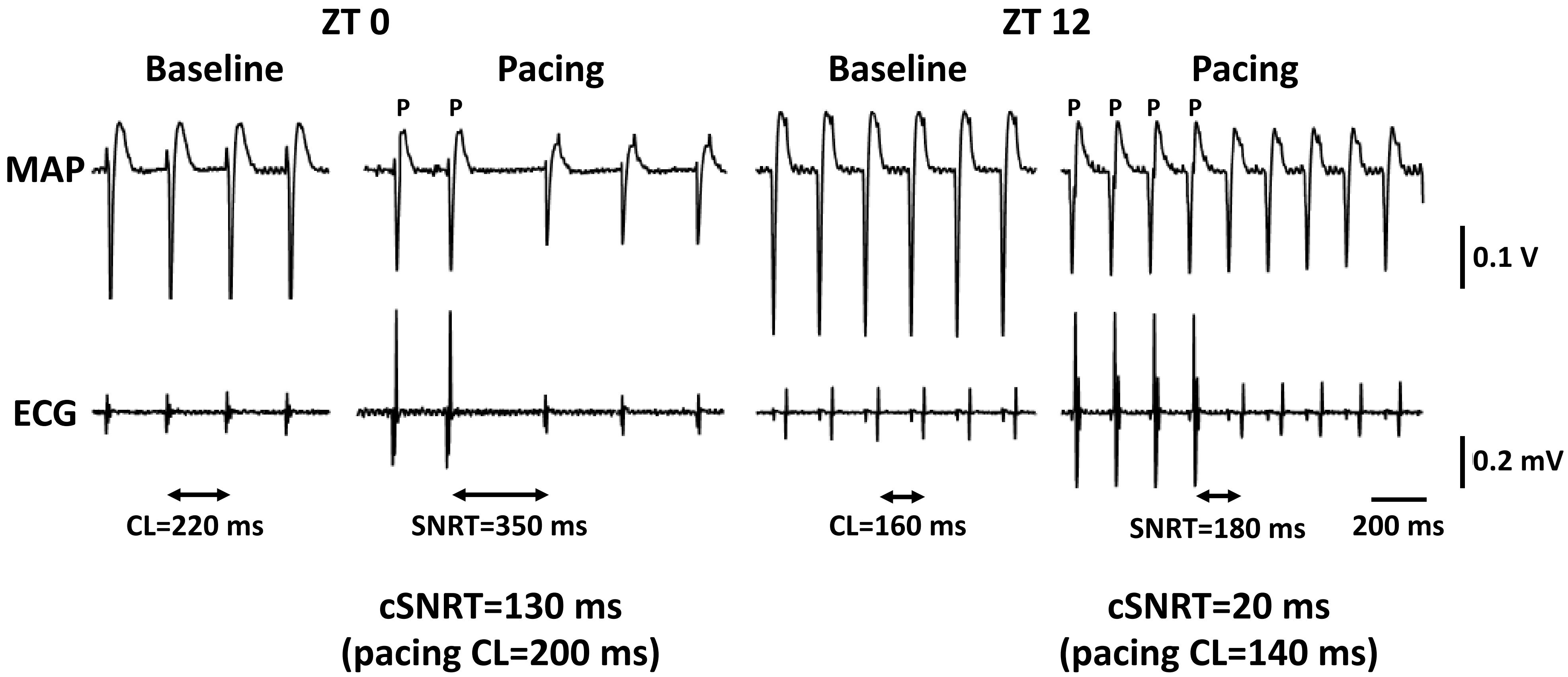


**Figure S1. Sinus node recovery time (SNRT) measurement.** Right atrial monophasic action potential (MAP) recordings and ECG-like recordings (using two electrodes placed on left atrium and ventricular apex) were made from Langendorff-perfused hearts from mice culled at ZT 0 and ZT 12. Recordings are shown under baseline conditions and at the end of 45 s pacing at a cycle length of the spontaneous cycle length minus 20 ms (P denotes paced beats).

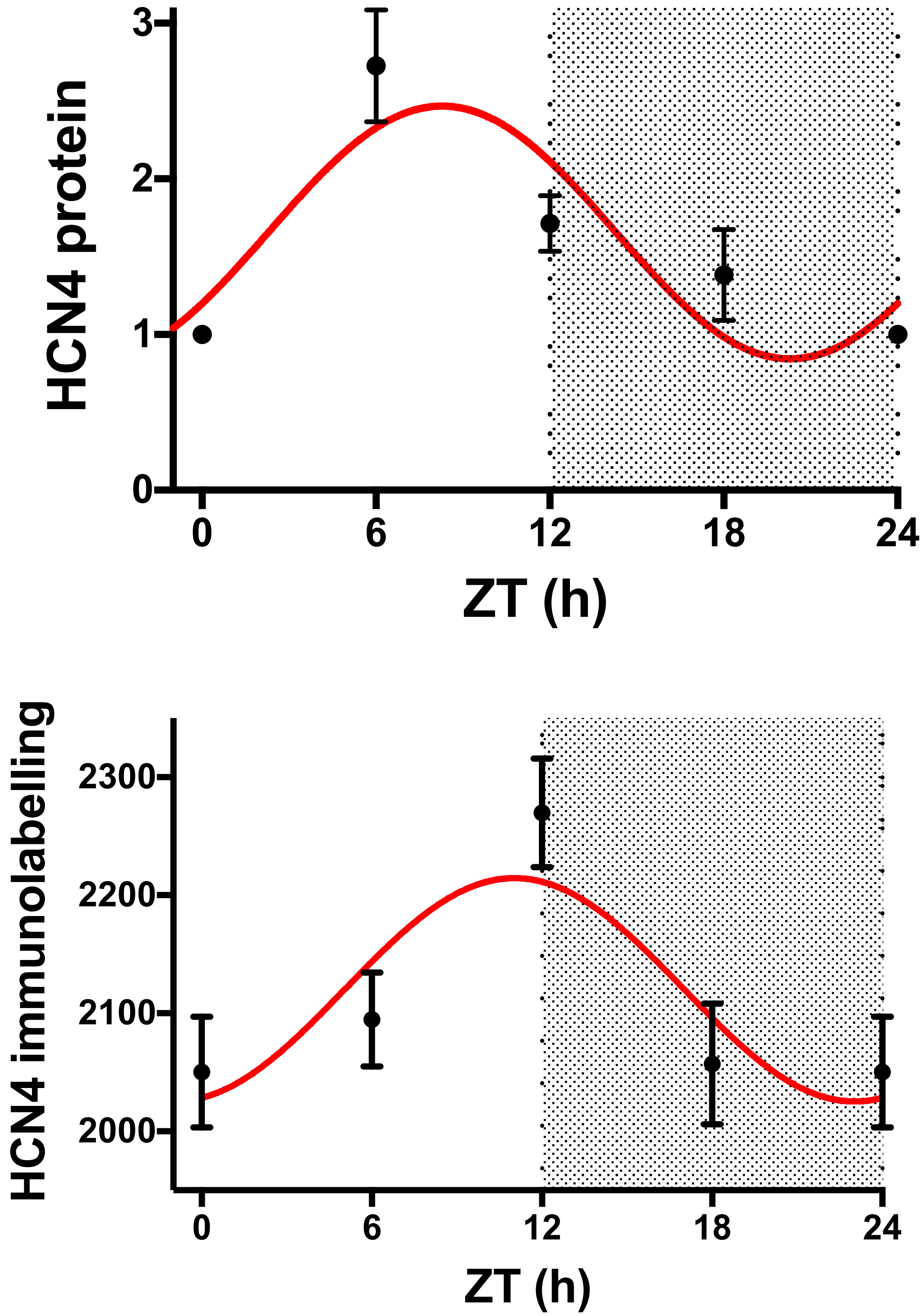


**Figure S2. Circadian rhythm of HCN4 protein expression. Top,** Expression of HCN4 protein as determined by western blot at four time points over 24 h (n=7 mice for ZT 0; n=4 for ZT 6; n=7 for ZT 12; n=4 for ZT 18; average of three technical replicates for each time point). **Bottom**, Expression of HCN4 protein as determined by immunohistochemistry at four time points over 24 h (n=3 mice for ZT 0; n=3 for ZT 6; n=3 for ZT 12; n=3 for ZT 18).

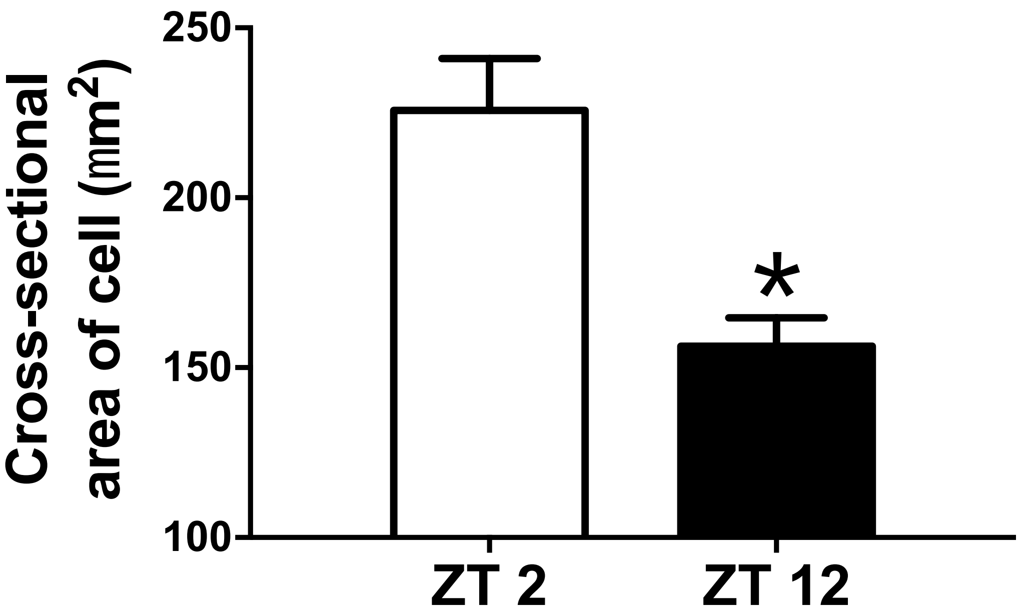


**Figure S3. Circadian rhythm in sinus node cell size.** The cross-sectional area of sinus node cells isolated at ZT 2 and ZT 12 was measured in Fluo 8-loaded cells (n=13 cells from 3 mice for ZT 0; n=13 cells from 3 mice for ZT 12. *P<0.05; two-tailed unpaired t test with Welch's correction. From the same experiments as Figures S4 and S5.

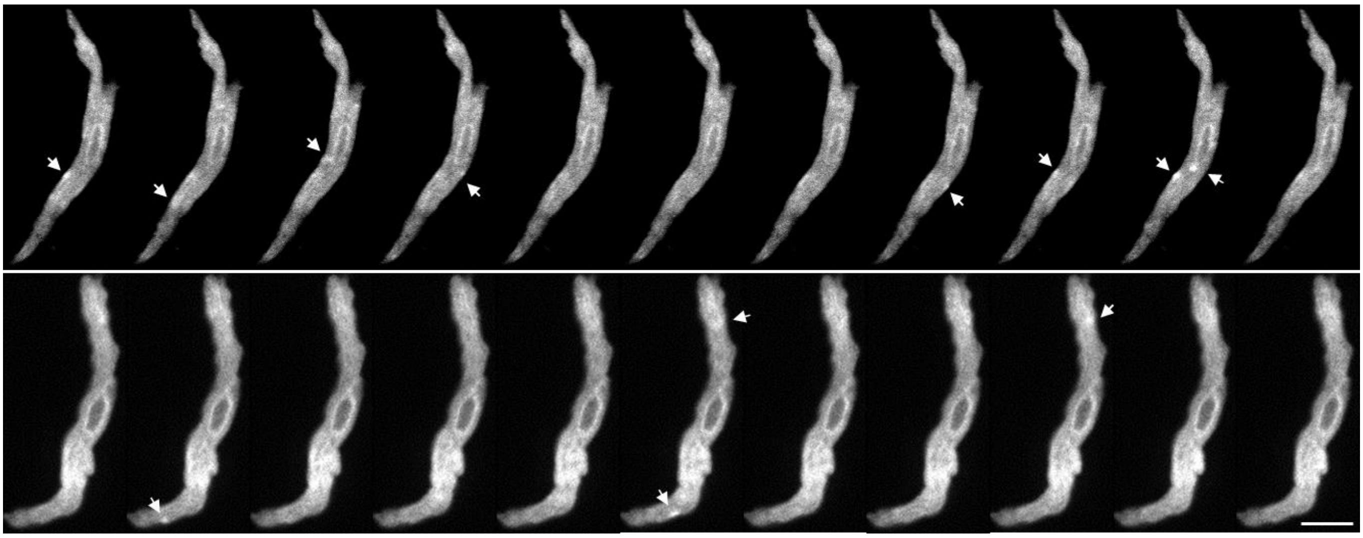

**Figure S4. Example of $Ca^{2+}$ sparks in an isolated sinus node cell.** The cell was loaded with Fluo 8 AM calcium indicator dye. The time interval between each image is 6 ms. White arrows indicate $Ca^{2+}$ sparks.

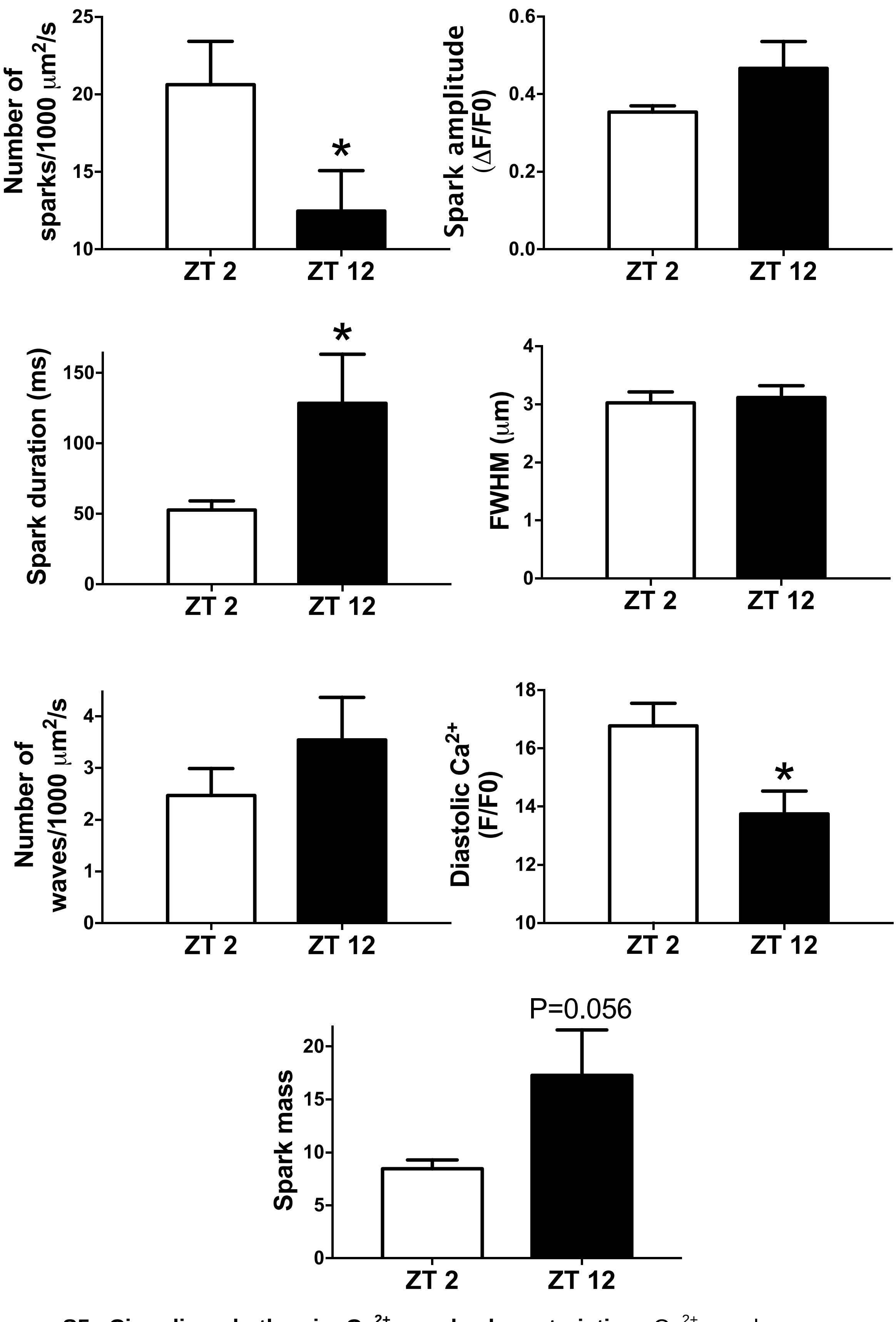


**Figure S5. Circadian rhythm in $Ca^{2+}$ spark characteristics.** $Ca^{2+}$ spark measurements are shown at ZT 2 (n=13 cells from 3 mice) and ZT 12 (n=13 cells from 3 mice). Number of sparks normalised to cell area and time, $Ca^{2+}$ spark amplitude, duration and FWHM (full width at half maximum amplitude of sparks), number of $Ca^{2+}$ waves normalised to cell area and time, diastolic $Ca^{2+}$ and spark mass shown. *$P<0.05$; number of sparks and waves tested by two-tailed Kolmogorov-Smirnov test and others tested by two-tailed unpaired t test with Welch's correction.

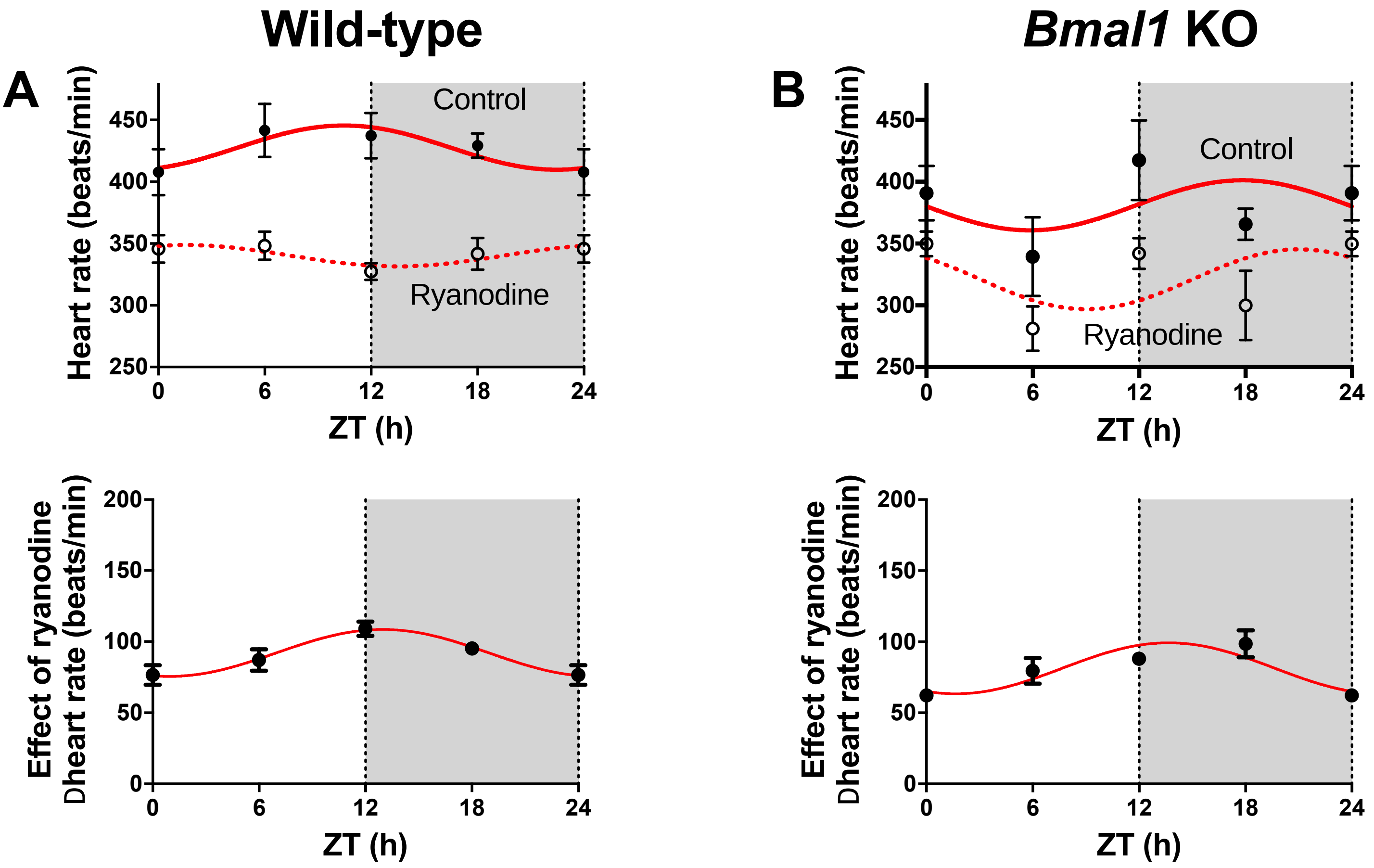


**Figure S6. Circadian rhythm in the $Ca^{2+}$ clock.** **A**, Intrinsic heart rate before and after the application of 2 µM ryanodine (top) and change in intrinsic heart rate after application of ryanodine (bottom) measured in the isolated sinus node from wild-type mice (n=11 mice for ZT 0; n=5 for ZT 6; n=9 for ZT 12; n=5 for ZT 18). **B**, Intrinsic heart rate before and after the application of 2 µM ryanodine (top) and change in intrinsic heart rate after application of ryanodine (bottom) measured in the isolated sinus node from cardiac-specific *Bmal1* knockout mice (n=8 mice for ZT 0; n=5 for ZT 6; n=5 for ZT 12; n=5 for ZT 18).

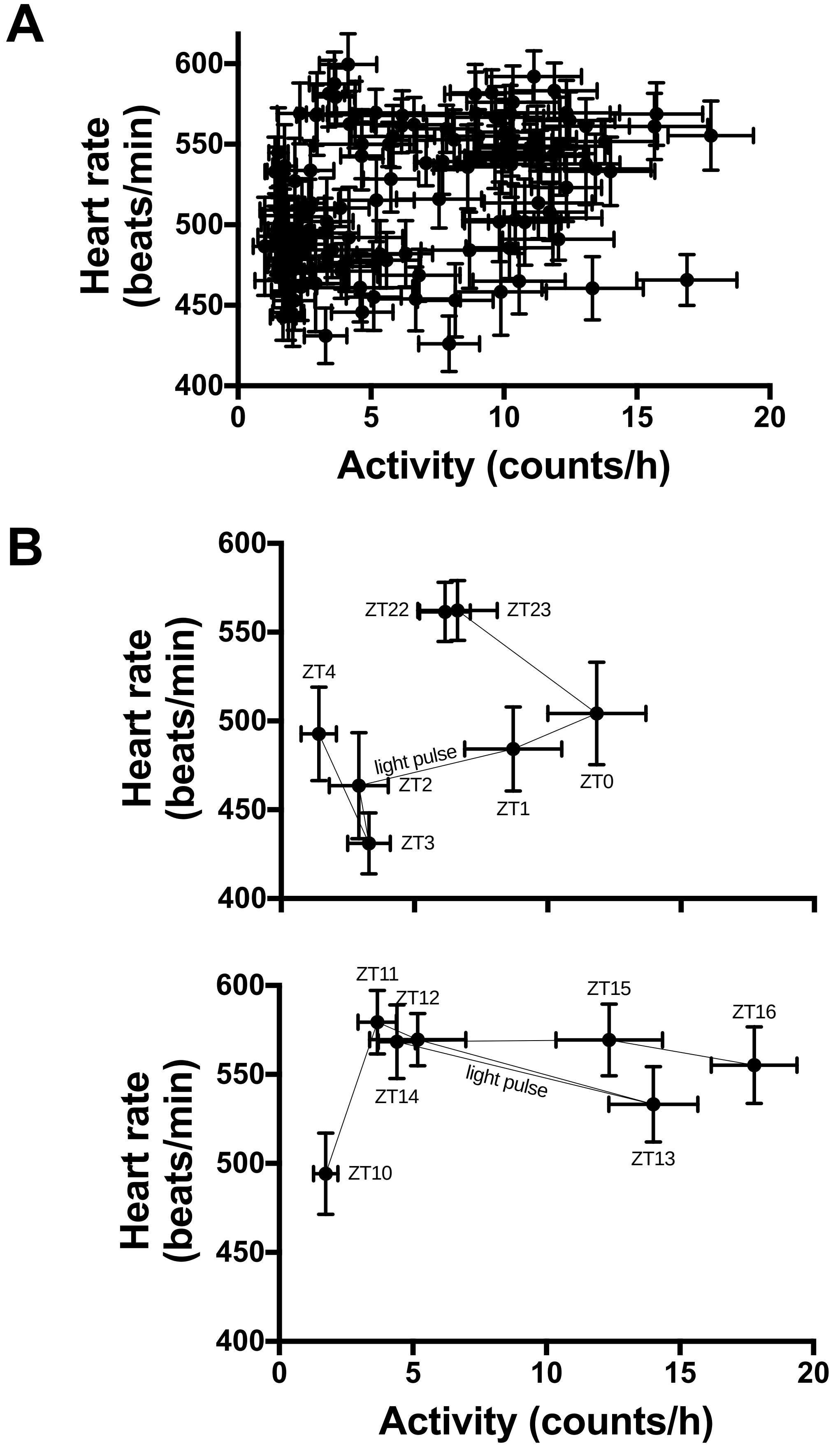


**Figure S7. No correlation between *in vivo* heart rate and physical activity.** The data shown are from the experiment shown in Figure 1. **A**, Relationship between *in vivo* heart rate and physical activity throughout the 6-day experiment shown in Figure 1. **B**, Relationship between *in vivo* heart rate and physical activity before, during and after the 1 h light pulses shown in Figure 1 and in more detail in Figure 7A.

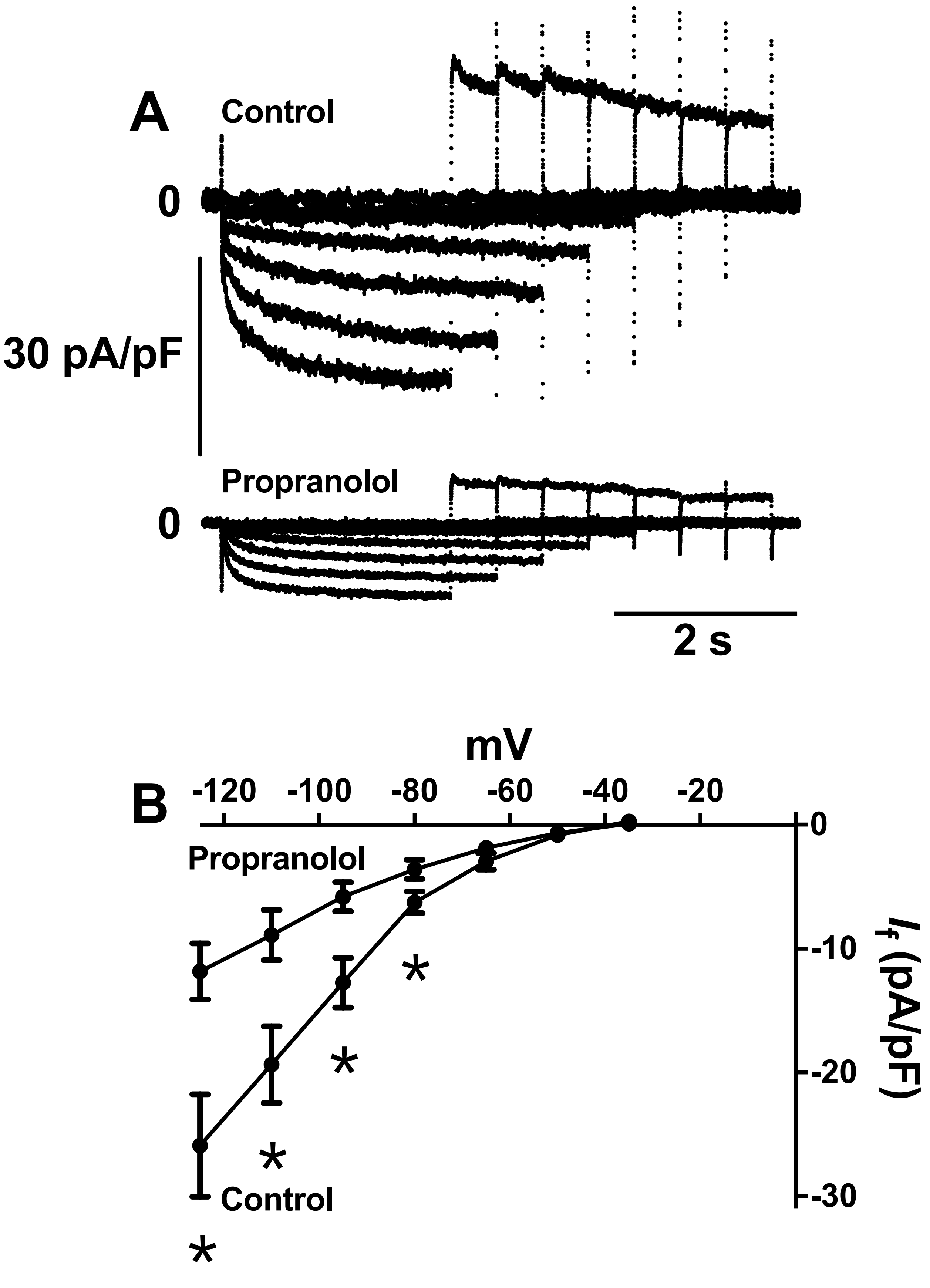


**Figure S8. Effect of 68.4 μM propranolol on $I_f$ in sinus node cells. A**, Families of recordings of $I_f$ in the absence (Control) and presence of propranolol. **B**, Current-voltage relationships for $I_f$ recorded in the absence and presence of propranolol (n=12 cells/3 mice). *P<0.05; two-way ANOVA with Dunnett's post-hoc test.
